# Population-scale subcellular proteomics reveals intracellular remodelling across the Alzheimer’s disease-resilience spectrum

**DOI:** 10.64898/2026.08.20.746004

**Authors:** Helen A. Jolly, Paula Seghers, Kaleah Balcomb, Amelia J. Smith, Laura Pearson, Ishan Agrawal, Bethany Geary, David A. Bennett, Thomas Wisniewski, Stephanie L. Fowler, Eleanor Drummond, Oliver M. Crook, Becky C. Carlyle

## Abstract

Proteome-wide analyses of human tissue have transformed our understanding of disease, but provide limited insight into protein localisation, a functionally informative dimension of the proteome. In Alzheimer’s disease, amyloid-β and tau exhibit aberrant localisation, yet whether spatial reorganisation extends proteome-wide has remained inaccessible to abundance-based proteomics. Here, we develop comparative subcellular proteomics applied to dorsolateral prefrontal cortex from 75 individuals spanning the Alzheimer’s disease-resilience spectrum, modelling protein localisation across disease. We identify 217 disease-associated localisation shifts enriched for endolysosomal function, intracellular trafficking, and RNA processing, and resolve tau proteoforms within insoluble aggregates. Our strongest localisation candidates show only modest differences in whole-tissue abundance, highlighting disease biology inaccessible to conventional proteomics. We validate co-localisation of CSNK1A1 with pathological tau and identify an unexpected neuronal localisation pattern for SCAI, a cancer-associated protein not previously characterised in human brain, highlighting the discovery potential of subcellular proteomics in tissue.

## Introduction

Cellular function is coupled to the spatial organisation of proteins, which governs their interactions, conformation, and access to signalling pathways(1–3). Disruption of intracellular organisation is a significant feature of Alzheimer’s disease (AD), whereby hallmark neuropathologies amyloid-β (Aβ) and tau exhibit mis-localisation, loss of native function, and neurotoxic prion-like aggregation. The accumulation and spread of these pathologies correlate with synaptic loss, neuroinflammation, and progressive cognitive decline(4–8). Cognitively resilient individuals have preserved cognitive function despite substantial neuropathological burden(9–11). This divergence between pathology and clinical phenotype suggests protective cellular mechanisms that are not fully characterised.

High-throughput proteomic studies of postmortem brain tissue have broadened understanding of the molecular landscape of AD beyond amyloid-β and tau, uncovering widespread disruption of synaptic, vascular, neuroinflammatory, and metabolic pathways(12–17). In bulk tissue studies, however, cytoarchitectural changes such as synapse loss and glial recruitment can confound protein abundance quantification, such that observed differences partly reflect shifts in tissue composition. Protein localisation offers a complementary and mechanistically informative dimension to the proteome, orthogonal to bulk expression(18).

Subcellular proteomics enables high-throughput interrogation of the spatial proteome. Established experimental methods, such as localisation of organelle proteins by isotope tagging (LOPIT)(19–21), dynamic organellar mapping (DOM)(22–24), and SubCellBarCode(25), separate tissue or cell homogenate on a biochemical or density gradient, such that proteins co-fractionate according to the physical properties of their subcellular niche. Mass spectrometry-based quantification of each fraction yields characteristic protein fractionation profiles that support inference of steady-state localisation, through protein correlation profiling(26–29).

Collectively, these approaches have demonstrated that protein localisation can be resolved at proteome scale based on fractionation behaviour across diverse *in vitro* systems, including those with complex cellular architecture(21,30,31). Beyond static maps, subcellular proteomics has enabled the detection of condition-dependent relocalisation events(22–24,32,33), and contributed to the characterisation of distinct subcellular niches(34). The application of machine learning frameworks has advanced with increasingly high throughput experimental data, enabling robust localisation inference(35,36), including probabilistic Bayesian approaches that capture multi-localisation and uncertainty(37–39).

Application of subcellular proteomics to multicellular tissue is emerging, with recent studies demonstrating feasibility in rat liver(40) and mouse heart(41). In AD brain, fractionation-based approaches have largely been applied as a compartment enrichment strategy, targeting membranes, synaptosomes, and extracellular vesicles(42–47). Kandigian *et al.* recently demonstrated that reproducible differential ultracentrifugation (DC) based fractionation is achievable in a small series (n = 5) of control and AD angular gyrus samples(18), establishing proof-of-principle that spatial proteomic organisation can be resolved in postmortem brain tissue despite the challenges associated with disease heterogeneity, postmortem interval, and tissue freezing(48,49). However, implementing this approach with larger sample sizes, required to interrogate inter-individual variability across the AD spectrum, has not been previously attempted.

While bulk proteomic studies in AD have increased to hundreds of samples(14,50), extending the scale of subcellular approaches presents an opportunity to systematically interrogate protein localisation dynamics across the disease spectrum, while considering inter-individual variation in neuropathology and clinical presentation.

Here, we establish subcellular organisation as a quantitative molecular phenotype at cohort-scale, using dorsolateral prefrontal cortex from 75 individuals spanning the AD-resilience spectrum in the ROSMAP cohort(51). Tissue was separated into seven subcellular fractions by differential ultracentrifugation and profiled using data-independent acquisition tandem-MS (DIA LC-MS^2^). We have adapted machine learning frameworks previously applied to cell-based systems(38) to model protein localisation in heterogeneous tissue, integrating clinical and neuropathological covariates to quantify disease-associated shifts in steady-state localisation. Rather than treating fractionation as a route to a single consensus map, this reframes localisation as a feature that varies amongst individuals, allowing subcellular reorganisation to be modelled as a function of disease classification, pathology, and resilience.

This approach highlights coordinated reorganisation across multiple subcellular systems. Differentially localised proteins (n = 217) are enriched for vesicle-mediated transport, endolysosomal function, and axonal trafficking. Concordant altered localisation patterns are observed for multiple components of defined molecular systems, including the AP-3 adaptor complex, the dynactin-mediated transport machinery, regulators of phosphoinositide metabolism, and the U1-spliceosome, pointing to convergent perturbation of intracellular trafficking and RNA processing dysfunction. Tau peptides derived from seed-competent domains display the greatest localisation divergence, demonstrating that our framework resolves disease-relevant proteoforms for a single protein.

While our findings capture established features of AD biology, we identify candidates that have not been prioritised in previous global abundance analyses. Our top result, protein SCAI (suppression of cancer cell invasion) has not been characterised in human brain, despite high expression. Tissue enrichment strategies, immunohistochemistry, and immunofluorescent imaging in cortical i^3^Neurons support a distinct localisation pattern in human brain, demonstrating the discovery potential of tissue-based subcellular methods.

### ROSMAP cohort

The Religious Orders Study (ROS) and Rush Memory and Aging Project (MAP) are community-based cohort studies of aging and dementia. Participants enrol without known dementia and agree to annual clinical evaluation and brain donation. Both studies were approved by an Institutional Review Board of Rush University Medical Center. All participants signed an informed and repository consent and Anatomic Gift Act. ROS includes older Catholic nuns, priests, and brothers from across the USA; MAP includes older persons from northeastern Illinois(51). All participants were classified at each annual visit for dementia and its causes, for example, Alzheimer’s dementia and mild cognitive impairment (MCI); participants without dementia or MCI were labelled no cognitive impairment (NCI) as previously reported (52–54). At the time of death, a neurologist reviewed select information from all visits and rendered a final summary diagnosis blinded to all pathologic data. A complete autopsy was done including Braak Stage, CERAD (consortium to establish a registry for AD), and ADNC (AD neuropathological change)(55). Based on a combined histopathological and clinical phenotype (**Table 1**), 75 ROSMAP participants (matched for Age, Sex, and PMI) were categorised for this study as control (CON, n = 25), cognitively resilient (RES, n = 25), and AD-dementia (AD-DEM, n = 25). Additional covariate details are shown in **Figure 1A**.

**Figure 1.**
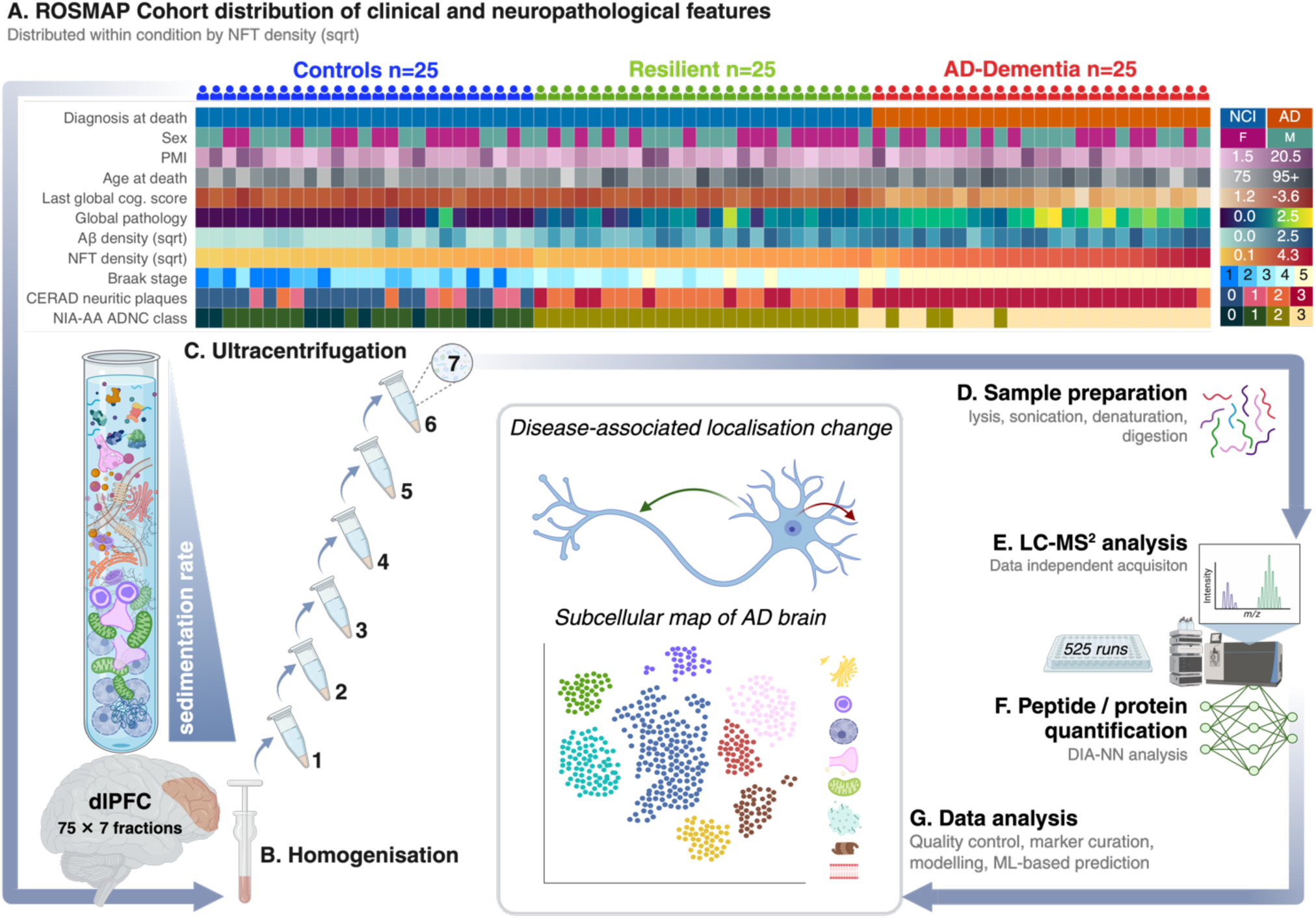
A) Distribution of both continuous and categorical ROSMAP covariates within disease class highlights intraclass variability. Subjects, classed as CON, RES, and AD-DEM, are ordered within-class by the square-root transform of their neurofibrillary tangle (NFT) density(51,53,56–58). Global cognition score was obtained from the average of 19 cognitive tests as described previously(61). Global pathology was quantified as the standard deviation-scaled count of both neuritic and diffuse plaques, and neurofibrillary tangles (NFTs), averaged across five brain regions(62). Density scores represent the proportion of Aβ and NFT positive regions calculated from 20-micron sections across eight regions, square-root (sqrt) transformed (63). PMI; postmortem interval in hours, CERAD; Consortium to establish a registry for AD, NIA-AA; National Institute on Aging and Alzheimer’s Association, ADNC; AD neuropathologic change. The right-hand legend shows: NCI; no cognitive impairment, AD; diagnosis of Alzheimer’s disease at death, F; female, M; male. For sqrt-transformed pathology variables, a value of 0.0 indicates absence. CERAD neuritic plaque score indicates their presence per mm^2^: 0; none, 1; sparse, 2; moderate, 3; frequent. NIA-AA-ADNC class represents likelihood of AD determined postmortem: 0; none, 1; low, 2; intermediate, 3; high. B,C) Differential ultracentrifugation-based subcellular fractionation, applied to all 75 dlPFC samples. D–F) Proteomic quantification of all 525 sample-fractions by DIA LC-MS^2^. G) Protein localisation prediction and inference of differential localisation.

**Table 1.** Clinical diagnoses reflect the final summary diagnosis (blinded to postmortem data) determined from review of select information from each annual evaluation, coded NCI (no cognitive impairment) and AD (Alzheimer’s-type dementia with no other cause of dementia) (56). Braak stages 1–2 indicate NFTs are confined to transentorhinal region, while stages 3–4 and 5–6 indicate spread across limbic and neocortical regions, respectively(53). Braak 3+ is required, but not sufficient to cause clinical AD. At stage 3, CON and RES individuals are distinguished by CERAD score and AD neuropathologic change (ADNC) class(57,58).

| Condition | Abbreviation | Definition<br>(NCI, no cognitive impairment; AD,<br>Alzheimer's disease) | Braak<br>Stage | Clinical<br>Score | CERAD<br>score | ADNC<br>class |
| --- | --- | --- | --- | --- | --- | --- |
| Healthy control | CON | NCI with low pathology | 1–3 | NCI | Inter-high | A |
| Cognitively resilient | RES | NCI with high pathology | 3–5 | NCI | Non-low | C |
| Alzheimer's disease | AD-DEM | Clinical AD with high pathology | 4–5 | AD-dementia | None-low | C |

## Results

### Study summary

A full summary of the ROSMAP participant (Subject) characteristics (**Figure 1A**) highlights the distribution of neuropathological and clinical covariates. Subjects are classed as controls (CON), resilient (RES), or AD-dementia (AD-DEM), as described in **Methods**. For sample processing, dorsolateral prefrontal cortex (dlPFC) tissue samples (n = 75) were homogenised and separated into seven fractions by serial ultracentrifugation (**Figure 1B, C, Supplementary figure S1**)(18,22) (**Methods, Supplementary SOP001**). DIA LC-MS^2^ followed by DIA-NN 2.0(59) identification (**Figure 1D–F**) yielded a full matrix of 118,740 precursor (peptide ions) mapped to 102,902 unique peptides, and a full protein-level matrix of 8,525 unique UniProt(60) IDs mapped to 8,510 parent gene names. Of these, 78,676 precursors from 70,118 unique peptides, and 7,271 unique proteins mapped to 7,264 gene symbols were retained for downstream analyses after quality-control filtering (**Methods**).

### Biochemical fractionation reproducibly resolves subcellular structure in postmortem brain tissue

First, we asked whether biochemical fractionation of postmortem brain tissue retains sufficient spatial information to distinguish subcellular organisation across individuals. Protein fractionation profile (sum normalised and z-scaled mean) clusters (Ward’s D2, k = 15)(64) are significantly enriched (GO:CC)(65,66) for distinct subcellular compartments, highlighting separation of both major membrane-bound structures and cytosolic intracellular components (**Figure 2A**). Consistent with this, 782 curated markers representing 11 subcellular compartments (**Figure 2B**) exhibit characteristic fractionation profiles (**Figure 2Ci, ii, Supplementary Figure S2**), and cluster according to their canonical localisation (**Figure 2Ciii**). Nuclear, mitochondrial, ribosomal, and peroxisomal markers show clearer separation, whereas plasma membrane, secretory pathway, and cytoskeletal proteins show greater overlap, representing compartments for which biochemical separation provides lower spatial resolution. Notably, protein fractionation profiles are highly correlated between individuals across the three disease groups (**Figure 2D**), demonstrating that subcellular fractionation can be applied at scale to reproducibly capture spatial proteomic information in brain tissue.

**Figure 2.**
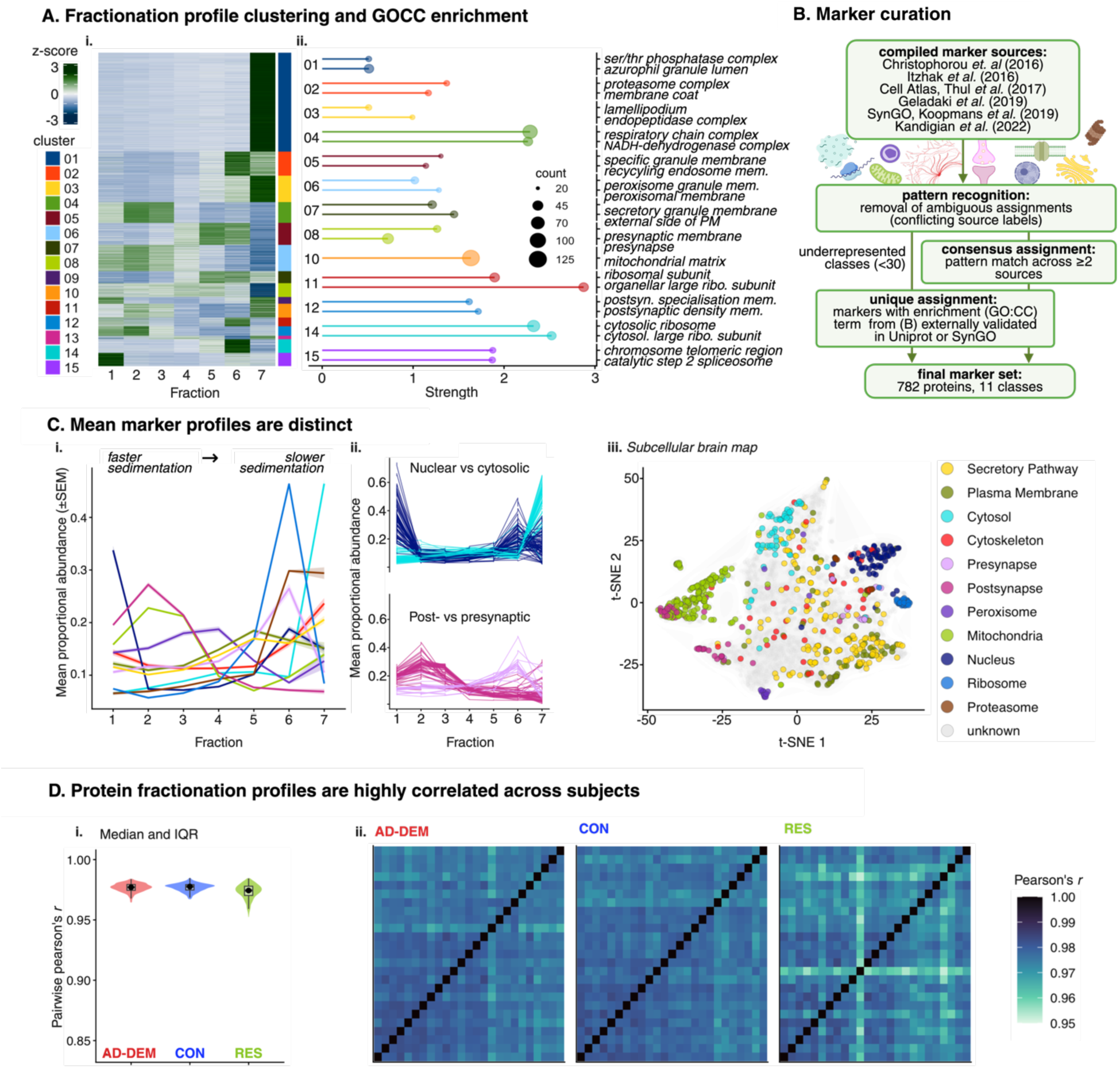
A) Heatmap (elements are coloured by z-score, shown in key) of mean sum-normalised and z-scaled protein intensities by fraction (x-axis) grouped by hierarchical clustering (Ward’s D2)(67) with k = 15 clusters. For example, cluster 01 represents proteins with the highest relative abundance in fraction 7, due to slower sedimentation rate. Cluster membership is indicated by colour (also in legend). Gene Ontology Cellular Component (GO:CC)(65,66) enrichment of each k cluster is shown in the right panel, with strength (log10-fold enrichment) on the x-axis, and GO:CC term on the y-axis. The number of protein entries overlapping with that term (count) is indicated by the lollipop size. B) Marker curation workflow, summarising the integration of different experimental subcellular proteomics marker sets to identify assignments that are the most robust among studies. C) Ribbon plot (i) shows the compartment class mean of proportional marker profiles. Standard error of the mean (SEM) is represented by ribbon width. For example, mean (across subjects) sum-normalised profiles of individual marker proteins: nuclear against cytosolic, and presynaptic against postsynaptic, show distinct proportional intensity peaks for these classes (ii). T-distributed stochastic neighbour embedding (t-SNE)(68) of sum-normalised and scaled mean fractionation profiles shows clustering of proteins by marker class (iii). D) Subject-by-subject pairwise correlations of log2-transformed fractionation profiles are stably high (Pearson’s r = 0.952– 0.985) across disease groups, which indicates reproducible quantification of subcellular proteomic signal across individuals. Violin plot (i) and heatmaps (ii) show distribution and pattern of correlations, respectively.

### Coordinated intracellular reorganisation of the subcellular proteome associates with AD

Given the reproducibility of proteomic fractionation profiles across individuals, we hypothesised that disease-associated subcellular reorganisation would be quantifiable as robust differences in protein fractionation between groups.

To determine the sources of variation that should be accounted for when modelling these differences, we first decomposed protein-level variance across biological and technical variables. Fraction identity accounts for the largest proportion of variance, consistent with the spatial information captured by biochemical fractionation. Disease condition (CON, RES, AD-DEM) also contributes substantially; proteins such as APP, NPTX2, SMOC1, and VGF are among the highest contributors to disease-associated variance, consistent with previous abundance-based proteomic studies(14) (**Figure 3A**).

**Figure 3.**
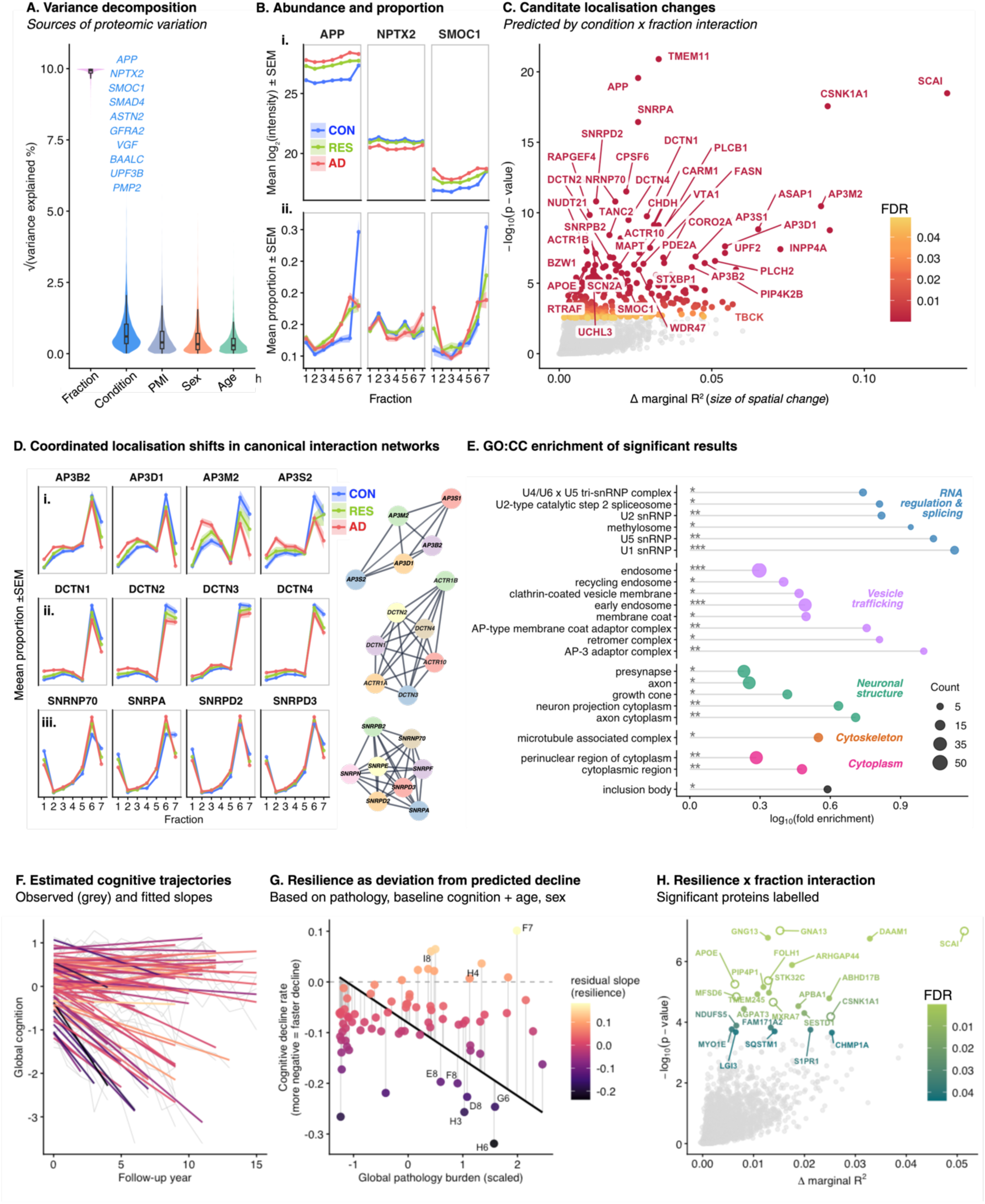
A) Variance decomposition for all fixed effects (x-axis) included in the interaction model showing labelled proteins that contribute to the largest amount of square-root scaled variance in log2-scaled protein intensities (y-axis). Proteins are labelled in order of variance contribution. Condition (CON, RES, and AD-DEM), PMI; postmortem interval, Age; age at death. B) The mean log2-scaled abundance and sum-normalised fractionation profiles, for APP, NPTX2, and SMOC1, shaded by condition. Ribbon width represents +/− standard error of the mean (SEM). The top panel shows relative abundance differences by disease, which highlights abundance differences consistent with traditional bulk proteomics. The bottom panel highlights the mean proportion of a protein found in a given fraction, sum-normalised to control for inter-individual differences in abundance. The larger peak in fraction 7 in CON is consistent with an increased soluble pool of proteins such as APP and SMOC1. In the case of APP, a higher peak for AD and RES in fraction 6 may represent the transition to(37) increased insoluble APP assemblies. C) Volcano plot of model results in unimputed dataset. Robust results (those replicated in analysis of both imputed and unimputed datasets, n = 217) are highlighted by FDR (false discovery rate). The proxy effect size for spatial differences (marginal R2 explained by the Condition × Fraction interaction term) is shown on the x-axis, against raw −log10(p-value) (y-axis). D) Subunits from each of the following complexes show concordant fractionation shifts by disease: AP3 adaptor (i), Dynactin (ii), and U1 spliceosome (iii), suggesting that differential localisation may be occurring at the level of large protein assemblies. Interaction networks of all FDR significant proteins with similar profile differences are shown(69,70) E) GO:CC(65,66) Log10-scaled fold enrichment (x-axis) of stable results (n = 217) highlights cellular components (y-axis) these proteins are enriched for. Asterisks indicate significance of observed enrichment; p < 0.05*, p < 0.01**, p < 0.001*** F) Observed global cognitive scores (grey lines) across longitudinal follow-up (x-axis) and subject-specific fitted trajectories (coloured), which are coloured by residual cognitive decline rate (resilience score). Warmer colours indicate subjects with slower-than-expected cognitive decline relative to pathology. G) Relationship between global pathology burden (x-axis) and model-estimated cognitive decline. The black line indicates the expected decline predicted by pathology, age at baseline, sex, and baseline cognition, while coloured points represent each subject’s estimated decline. Vertical segments show the residual deviation from predicted decline (resilience score), with positive residuals indicating greater cognitive resilience and negative residuals indicating greater vulnerability. Selected high-pathology resilient (I8, H4, F7) and vulnerable individuals (for example H6 and H3) are labelled. H) Significant (5% FDR) proteins from the resilience interaction model (unimputed dataset) are highlighted in the volcano plot. Stable results from analyses of both imputed and unimputed datasets are represented by empty circles.

We therefore modelled disease-associated differences in fractionation through the Condition × Fraction interaction (which captures how fractionation profile shapes differ between groups, **Figure 3B**), while controlling for overall abundance and relevant sources of inter-individual variation, including age at death, sex, and postmortem interval (**Methods**). This marker-independent strategy bypasses prior compartment assignment in multicellular human brain tissue, where reliable compartment-specific markers are not well defined and many proteins occupy multiple intracellular locations(71). Thus, disease-associated localisation differences can be detected even when proteins show no difference in global abundance or cannot be confidently assigned to a single subcellular compartment.

Across 4,794 proteins passing quality-control intensity and missingness thresholds, 236 and 285 proteins exhibit significant Condition × Fraction interactions in the imputed and unimputed datasets, respectively (FDR < 5%)(72). To minimise dependence on missing-data handling, subsequent analyses focus on the intersection of these results – a conservative set of 217 proteins showing robust disease-associated differences in subcellular distribution (**Figure 3C**). This set includes numerous proteins previously associated with neurodegeneration, including APP(73,74), APOE(75,76), MAPT(77,78), SMOC1(79), HTT(80), SQSTM1(81,82), STXBP1(83), S100A9(84,85), YWHAG/B(86), DCTN1(87), EIF4G1 and VPS35(88), and ASAP1(89), indicating that established disease biology extends beyond differential abundance to include altered subcellular organisation (**Figure 4, Supplementary Data A**). Notably, candidates with higher-ranking effect sizes (**Figure 3C**) have not been prioritised through traditional abundance-based proteomics, such as SCAI, CSNK1A1, AP3M2(82), and INPP4A(90) (**Supplementary Figure S3**).

**Figure 4.**
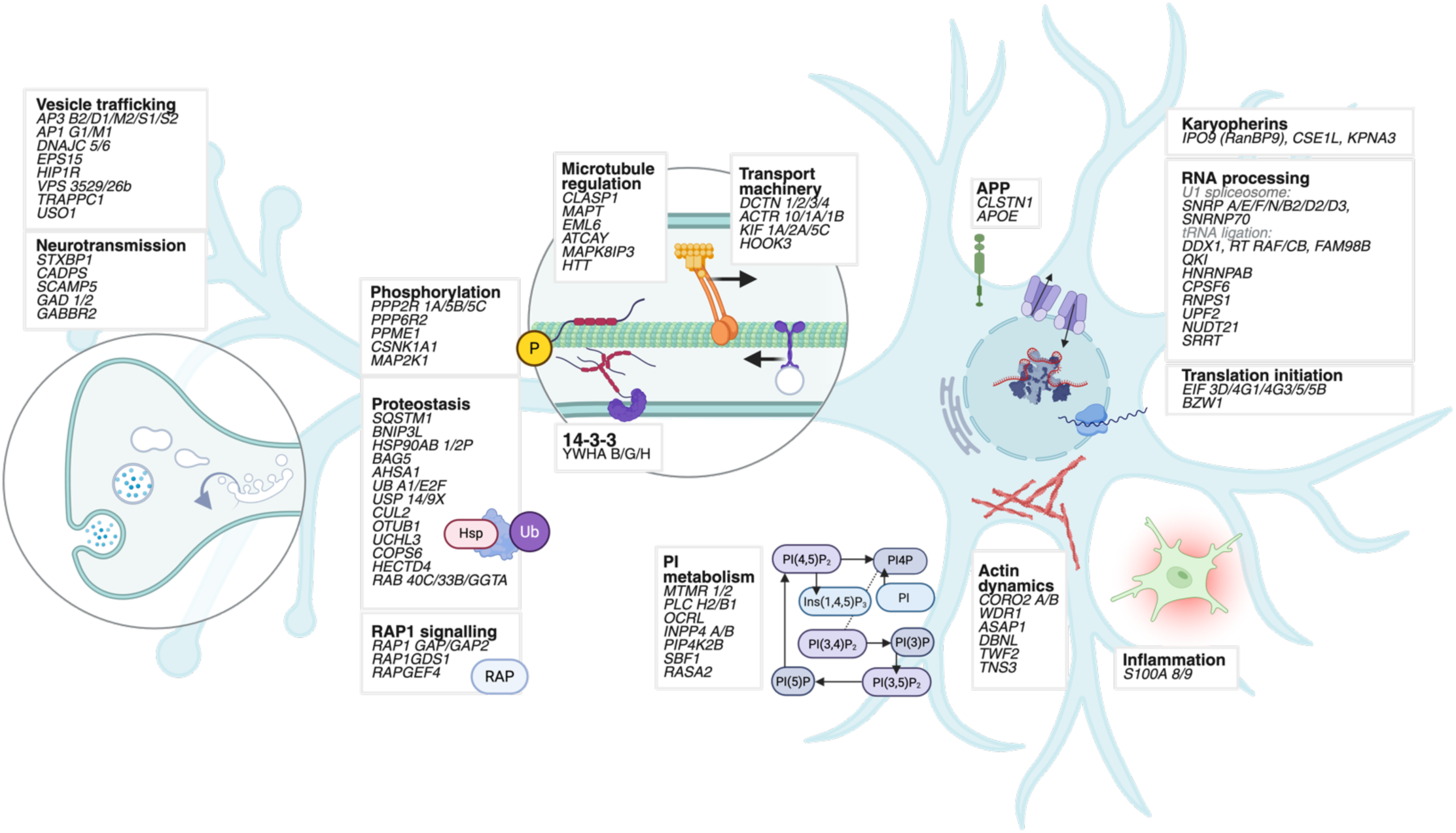
Schematic summary of the key biological processes exhibiting differential protein localisation in the categorical model (CON, RES, and AD-DEM). Candidate proteins are grouped according to their predominant cellular functions. Proteins within a box share at least one functional or physical association in STRING(69).

Resilient individuals exhibit neuropathological burden intermediate between controls and AD-DEM (**Figure 1A**), illustrating that pathology exists on a continuum rather than as discrete disease categories. We therefore also modelled localisation as a function of continuous neuropathological burden, replacing the categorical disease variable with a global neuropathology score summarised across five brain regions(62) (z-scaled; **Figure 1A, Methods**). This identifies 408 proteins with significant pathology × Fraction interactions in both the imputed and unimputed datasets (FDR < 5%) (**Supplementary Data B**). Of those also identified in the initial categorical analysis (n = 174/217), effect size estimates are highly correlated between models (Pearson’s *r* = 0.96, *p* < 2e^−16^). Top candidates, including SCAI and CSNK1A1, are recovered by both approaches, indicating that results are robust to disease parameterisation (**Supplementary Figure S3**).

#### Protein redistribution occurs through coordinated remodelling of localisation-dependent molecular systems

If disease-associated localisation differences are not randomly distributed across the proteome, they should instead converge on interacting proteins and connected cellular processes. STRING protein-protein interaction analysis(69) against a background of all modelled proteins demonstrates significant network enrichment (average local clustering coefficient = 0.43, *p* = 2.6×10^−10^), highlighting that proteins undergoing redistribution may form coordinated functional modules rather than isolated localisation events. Consistent with this interpretation, multiple subunits of canonical protein complexes exhibit concordant disease-associated fractionation profiles (**Figure 3D**). Examples include the dynactin complex (DCTN1–4, ACTR1A/B and ACTR10), the AP-3 adaptor complex (AP3D1, AP3B2, AP3M2, AP3S1 and AP3S2), the cargo-selective retromer (VPS29, VPS35 and VPS26B), the tRNA ligase complex (DDX1, RTRAF, RTCB and FAM98B), 14-3-3 proteins (YWHAH, YWHAB and YWHAG), and eight components of the U1 spliceosome (**Figure 3C**, **Figure 4, Supplementary Figure S4**).

Enrichment analysis (GO:CC)(65,66) similarly highlights protein systems involved in vesicle-mediated transport and axonal trafficking (**Figure 3E**), indicating that disease-associated redistribution may preferentially affect localisation-dependent molecular assemblies rather than individual proteins. Although not significantly enriched by ontology analysis, multiple components of the phosphoinositide (PI) signalling pathway, including INPP4A/B, PLCB1, PLCH2, OCRL, MTMR1/2, PIP4K2B and SBF1, reflect similar disease-associated change in fractionation profiles, characterised by a relative increase in earlier fractions and depletion from later fractions in AD (**Figure 4, Supplementary Figure S4**). These proteins represent distinct enzymatic steps within PI signalling pathways and have high-ranking effect sizes (ΔR² 2–7%). Together with related regulators such as ASAP1, TBCK, RAP1GAP, RAP1GAP2 and RELCH, these findings suggest that disease-associated spatial remodelling extends across interconnected pathways that coordinate membrane identity, vesicle trafficking and intracellular signalling.

#### Spatial correlates of cognitive resilience

Disease-group comparisons capture average differences between CON, RES, and AD-DEM, but collapse the substantial within-group heterogeneity that defines cognitive resilience: the capacity of some individuals to tolerate high neuropathological burden without commensurate cognitive decline. For example, SCAI and CSNK1A1 display highly similar fractionation profiles in CON and RES, whereas the principal redistribution occurs in AD-DEM (**Supplementary Figure S3**). We therefore modelled resilience as a continuous phenotype to identify localisation changes associated with individual rates of cognitive decline. We first estimated rates of cognitive decline using longitudinal global cognitive scores collected across study follow-up (644 observations from 75 individuals). The expected rate of cognitive decline was estimated for each individual, conditional on baseline cognition(91), baseline age, sex, and global neuropathological burden (**Figure 1A, Methods**). Subject-specific deviations from this expected trajectory (estimated as random slopes) were then used as a continuous measure of cognitive resilience, whereby slower-than-expected decline indicates greater resilience and faster-than-expected decline indicates greater vulnerability (**Figure 3F, Methods**).

Global cognition declines significantly over time (*β* = −0.081 per year, *t* = −5.07), while higher neuropathological burden is associated with faster decline (follow-up year × global pathology: *β* = −0.072, *t* = −5.44), consistent with expected patterns of disease progression. Baseline age also contributes to an increased rate of decline (*β* = −0.024, *t* = −2.01). Consistent with previous literature(91), higher baseline cognition is associated with a modestly slower rate of decline (*β* = 0.024, *t* = 1.77).

Using the resilience score (**Figure 3G**) as a continuous variable in our interaction model identifies 24 proteins at 5% FDR (**Figure 3H**, unimputed dataset), 7 of which are robust to imputation strategy (GNA13, APOE, SCAI, CSNK1A1, FOLH1, MXRA7, MFSD6). Though effect sizes are modest (ΔR² = 0.006– 0.051) relative to those observed for the categorical model, several proteins from the categorical analysis (CON, RES, AD-DEM) exhibit reproducible resilience-associated redistribution, including SCAI (ΔR² = 0.051), CSNK1A1 (ΔR² = 0.025), SQSTM1(ΔR² = 0.014), GNA13 (ΔR² = 0.015), and APOE (ΔR² = 0.006) (**Supplementary Data C**).

### Peptide-level analysis is sensitive to tau proteoforms

Tau (MAPT) represents a well-characterised example of proteoform-specific pathology in Alzheimer’s disease, therefore provides a useful benchmark for evaluating proteoform-level localisation using peptide-level data. The microtubule-binding repeat region (MTBR; R1–R4) contains the aggregation-prone PHF6* and PHF6 motifs that are sufficient to nucleate tau fibrillisation and are enriched within pathological tau assemblies. Consistent with this biology, individual tau peptides display marked heterogeneity in their fractionation profiles (**Figure 5**). Peptides derived from the N-terminal projection domain exhibit relatively stable fractionation patterns across disease groups. In contrast, peptides mapping to MTBR regions show pronounced disease-associated differences, characterised by greater proportional abundance within fractions 5 and 6 in AD subjects. In particular, the R2 domain shows the greatest disease divergence. The R2 repeat is exclusive to 4R tau, a more aggregation-prone proteoform containing both PHF6 and PHF6* nucleation motifs(92,93).

**Figure 5.**
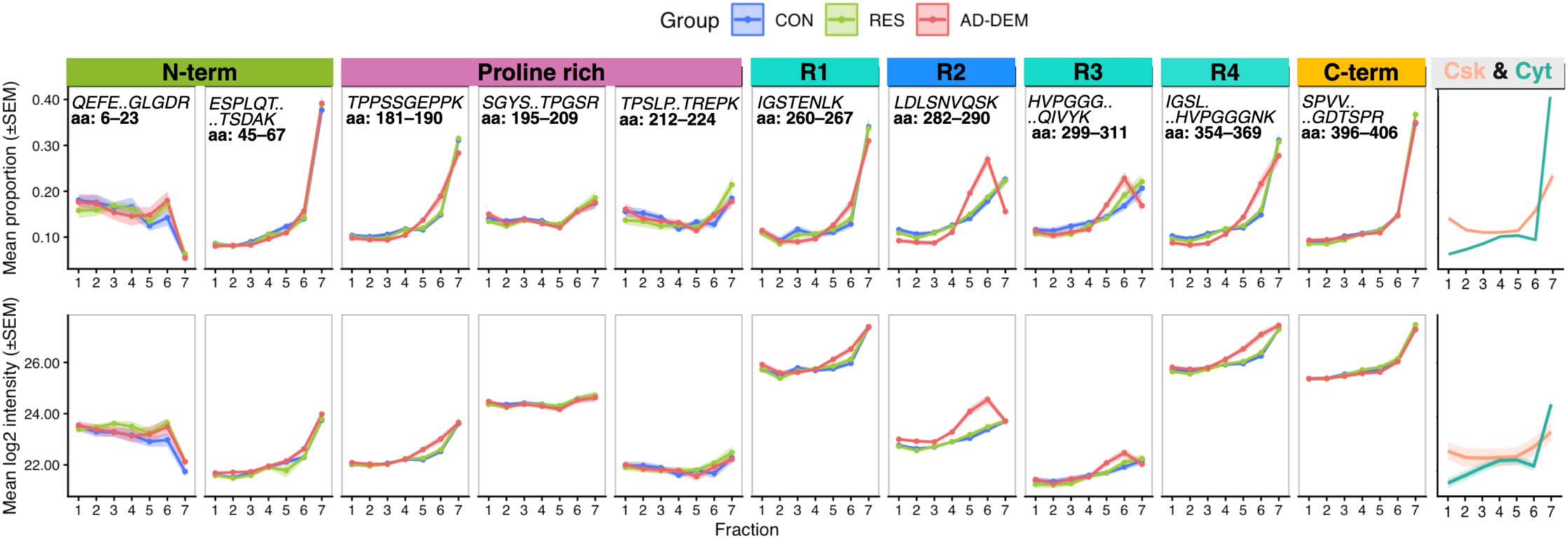
Domain-specific fractionation profiles of tau peptides (median precursor aggregated) across subcellular fractions. Upper panel (i) shows mean sum-normalised peptide abundance (y-axis), highlighting their proportional distribution across fractions (x-axis). Lower panel (ii) shows mean log2-scaled peptide intensity (y-axis) across fractions (x-axis), emphasising differential abundances. Ribbon width represents standard error of the mean (SEM). Peptides from the microtubule-binding repeat region display the greatest AD-DEM associated redistribution, while N-terminal peptides are comparatively stable. Mean cytoskeleton (Csk) and cytosolic (Cyt) marker profiles are shown on the right-hand panel for reference.

### Subcellular localisation can be predicted reproducibly across individuals

#### Localisation profiles generalise across individuals in out-of-sample validation

Having identified disease-associated changes in protein fractionation, we next asked whether subcellular localisation could be predicted reproducibly across individuals. We first generated individual subcellular maps for each subject (**Figure 6A**) using TAGM-MAP (T-Augmented Gaussian Mixture Model – Maximum A Posteriori), a machine-learning approach that uses 782 curated markers representing 11 subcellular compartments to learn their characteristic fractionation profiles (**Figure 2C**). For each protein, this provides a probability of localisation to each compartment based on how closely its fractionation behaviour resembles the corresponding marker proteins.

**Figure 6.**
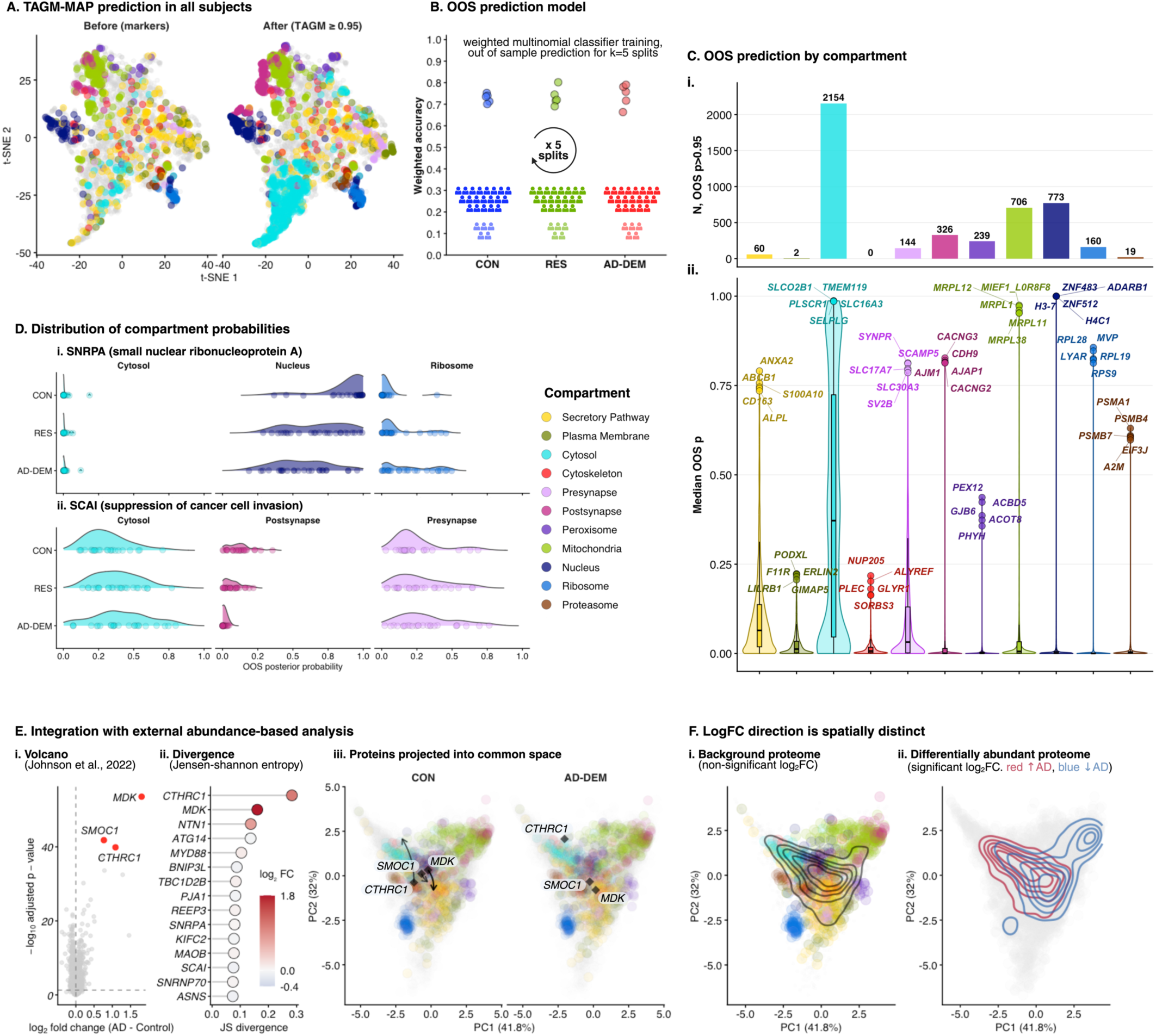
A) t-SNE of protein localisation before and after TAGM-MAP classification, highlighting confident (posterior probability × [1 - outlier probability] ≥ 0.95) assignment of proteins to 11 subcellular compartments. B) Accuracy of out-of-sample (OOS) localisation prediction framework, with schematic outlining subject-level five-fold cross-validation using weighted multinomial regression, trained independently within disease groups. C) The number of confidently assigned proteins (i) and the distribution of cohort median OOS localisation probabilities (ii) for all proteins by subcellular compartment. D) Logit-transformed OOS probability distributions for SNRPA and SCAI across diagnostic groups. E) Differential abundance from an independent bulk proteomic cohort (i) plotted against Jensen-Shannon divergence of OOS localisation distributions (ii), highlighting proteins exhibiting concomitant abundance and localisation differences. Projection of proteins in PCA space for each disease group (iii). All curated marker proteins are shaded by compartment to highlight spatial clustering. SMOC1; SPARC related modular calcium-binding protein 1, MDK; Midkine, CTHRC1; Collagen Triple Helix Repeat Containing 1. F) Density contours showing the distribution of the background proteome (top) and proteins with significant differential abundance in AD-DEM (bottom) in common PCA space. Increased and decreased abundance proteins are shown separately.

Across subjects, confident predictions (*p* ≥ 0.95) range from 499–1275 proteins per individual (**Supplementary Data D**). However, within-subject predictions do not quantify how reproducible protein localisation is across the cohort, or how well assignments generalise to unseen individuals. Consequently, we developed an out-of-sample (OOS) prediction model. For each disease group, this was trained on a subset of subjects and tested on held-out subjects (**Figure 6B**, **Methods**). In the OOS model, compartment classification accuracy is consistent across disease groups (**Figure 6B**, **Table 2**).

**Table 2.**
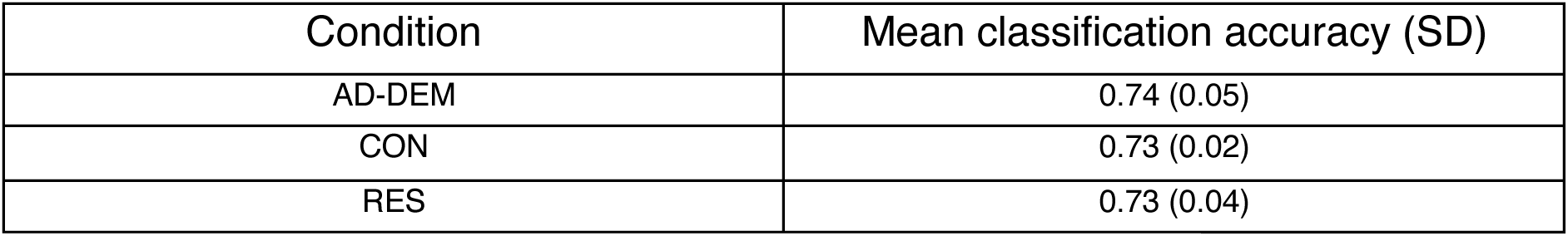
Accuracy and standard deviation (SD) of OOS prediction model.

| Condition | Mean classification accuracy (SD) |
| --- | --- |
| AD-DEM | 0.74 (0.05) |
| CON | 0.73 (0.02) |
| RES | 0.73 (0.04) |

Localisation is most consistently resolved for cytosolic, presynaptic, nuclear, mitochondrial, and peroxisomal compartments, with relatively few proteins reaching confidence (*p* ≥ 0.95) for the postsynapse, membrane, and secretory pathway (**Figure 6C**), likely due to limited resolution and the tendency for these compartments to co-fractionate, consistent with shared peaks in their marker distributions (**Figure 2C**).

Nonetheless, ranked set enrichment analysis (GO:CC)(66,94) (**Methods**, **Supplementary Data D**) demonstrates that the localisation model recovers biologically coherent subcellular organisation (**Supplementary Figure S5**).

Instead of requiring every protein to occupy a single subcellular compartment, this framework summarises localisation as a multi-compartment probability distribution, allowing differences in subcellular distribution to be visualised between disease groups. For example, the predicted localisation of SNRPA shifts away from the nucleus in AD-DEM (**Figure 6D, i**), consistent with its disease-associated fractionation change (**Figure 3D, iii**). By contrast, SCAI (**Figure 6D, ii**) has a broader localisation distribution that is more variable across RES and AD, consistent with localisation across multiple cellular compartments. In such cases, combining localisation prediction with our marker-independent analysis (**Figure 3**) identifies disease-associated differences in subcellular distribution even when proteins may be multi-localised within or across cell types, or cannot be confidently assigned to a single compartment.

#### Integration with large-scale bulk dataset highlights concomitant spatial and abundance difference

We then investigated how predicted changes in protein localisation relate to changes in overall protein abundance, identified in a proteomic analysis of bulk tissue from 110 control and 108 Alzheimer’s disease brains(14). Proteins with significantly different abundance in the external dataset (Holm adjusted *p* < 0.05) (**Figure 6E, i**) exhibit distinct fractionation patterns within our subcellular dataset (fractionation PCA space, **Figure 6E, iii**). Notably, the three proteins with the greatest abundance increase in AD (MDK, CTHRC1, and SMOC1) also have greater differences in localisation probabilities between disease groups (Jensen-Shannon divergence), from our OOS model (**Figure 6E, ii, Supplementary Data D**). Therefore, for some proteins, disease-associated abundance differences may occur alongside subcellular redistribution, representing an additional dimension of proteomic variation.

### SCAI is a cancer-associated protein with a putative role in neurons

Though differential localisation and abundance can coincide, our interaction model also identifies disease-associated localisation differences that may be less apparent to abundance-based proteomics. We selected SCAI and CSNK1A1 for orthogonal validation because both rank among the largest spatial effects (>5% marginal R²; **Figure 3C**), associate with resilience, yet show only subtle differences in whole-tissue abundance(14,15,17) (**Figure 7A**). SCAI is particularly unexplored, with enriched neuronal expression but no previous characterisation in human brain(95).

**Figure 7.**
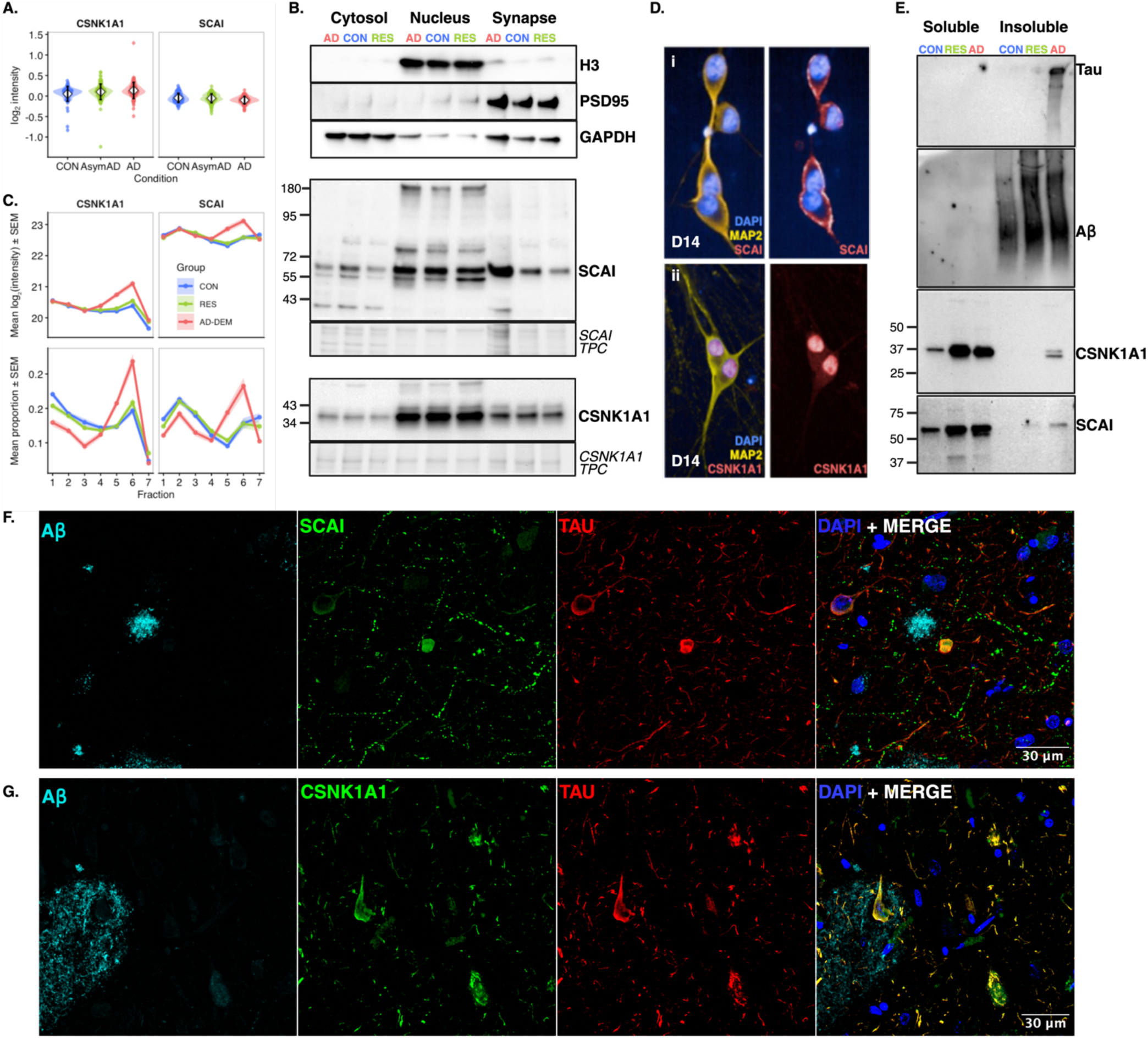
A) Modest abundance differences of CSNK1A1 (left) and SCAI (right) in the bulk abundance dataset from Johnson et al., 2022(14) by condition, classified in the study as controls (CON), asymptomatic AD (AsymAD), and AD. B) Western blot of SCAI and CSNK1A1 across nuclear, cytosolic, and synaptic-enriched fractions. Enrichment is validated by marker proteins H3 (nuclear), GAPDH (cytosolic), and PSD95 (synaptic), known to predominantly localise to these subcellular niches, respectively. Total protein content (TPC) is shown beneath for reference. The estimated mass in kDa is indicated on the left. C) Log2-scaled intensity (i) and proportional intensity (ii) for CSNK1A1 and SCAI highlight a peak in fractions 5 and 6 in AD, suggestive of a mass or density shift. Notably, profiles of CON and RES are similar. D) Immunocytochemistry (ICC) of SCAI (i) and CSNK1A1 (ii) in cortical i3Neurons at day 14 (D14), highlighting predominant membrane and nuclear association, respectively. E) Western blot of sarkosyl-soluble (left) and sarkosyl-insoluble (right) fractions of a CON, RES, and AD-DEM (AD) sample. Tau (AT8) is seen in the insoluble AD-DEM (AD) fraction but not in RES, while insoluble Aβ is detected with increasing intensity across CON, RES, and AD-DEM (AD) insoluble fractions. A double band is visible for CSNK1A1 (CK1a1) in the insoluble AD fraction. SCAI is also detected in the insoluble AD fraction. F) Immunohistochemistry (IHC) in sample AD2 cortex (Table 4, Methods) indicates an Aβ plaque and a tau-positive cell body. While present in this cell body, SCAI is not exclusively enriched within these cells across the cortex. Beaded processes are visible throughout the cortex, however, suggesting distinct localisation in brain. G) Immunohistochemistry (IHC) in sample AD2 cortex (Table 4, Methods). CSNK1A1 shows co-localisation with tau-positive neurons throughout the cortex in an AD sample.

To independently assess their subcellular distribution, dlPFC from a subset of the ROSMAP cohort (n = 3 per condition) was fractionated into cytosolic, nuclear, and synaptic-enriched compartments (**Methods**). SCAI is detected in nuclear, cytosolic and synaptic-enriched fractions (**Figure 7B, Supplementary Figure S8**), indicating broader multi-compartment localisation despite its predominantly nuclear annotation in existing databases(60,95). CSNK1A1 exhibits stronger nuclear enrichment with lower cytosolic signal, consistent with the localisation profile inferred from our cohort-scale fractionation analysis (**Figure 7C**). Although the low-throughput nature of these orthogonal analyses and restricted ability to normalise across the proteome limits robust quantification between disease groups, these findings independently support the subcellular distributions identified by biochemical fractionation, and highlight SCAI as a multi-localised protein in human brain.

SCAI (suppression of cancer cell invasion) has a defined role in cancer, where it acts as a negative regulator of MTRF/SRF transcription in the nucleus(96). Despite a proposed effect on dendritic morphology in rat neurons(97,98), SCAI has not previously been characterised in human neurons. Therefore, we next examined its localisation in cortical i^3^Neurons. SCAI exhibits prominent neuronal membrane localisation surrounding MAP2-positive cell bodies, with weaker cytoplasmic and nuclear staining (**Figure 7Di, Supplementary Figure S7**). CSNK1A1 has a predominantly nuclear pattern, with weaker cytosolic signal (**Figure 7Dii**).

Given the unexpected synaptic distribution of SCAI, we next examined whether its primary sequence contained features consistent with multi-compartment targeting. DeepLoc2.0(99) predicts a canonical N-terminal nuclear localisation signal. While DeepTMHMM1.0 (100) predicts a globular structure that is not membrane-spanning using the canonical residue sequence, UniProt annotates a hydrophobic C-terminal region (residues 472–492; GSLFTLFLNNPLMAFLFVSGLS) compatible with membrane association. This aligns more closely with our biochemical fractionation and immunocytochemistry, providing further support for multi-compartment localisation of SCAI in human brain.

Both SCAI and CSNK1A1 were markedly higher in fraction 6 specifically in AD-DEM, closely resembling the localisation profiles of APP and MTBR tau-derived peptides. Because these profile differences are consistent with a shift into insoluble species, we assessed their presence following sarkosyl extraction (n = 2 per condition). CSNK1A1 is detected within the insoluble fraction of AD-DEM cortex together with phospho-tau (AT8) and Aβ (6E10), while absent from control tissue (**Figure 7E, Supplementary Figure S9**). SCAI is visible in the insoluble fraction of an AD-DEM case but not in RES or CON, suggesting the presence of insoluble proteoforms (**Figure 7E, Supplementary Figure S9**). Notably, in a study of the ageing mouse brain proteome(101), SCAI displays the third highest increase in protein half-life in the hippocampus.

Multiplex immunofluorescence in samples from New York University Alzheimer’s Disease Center (**Methods**) (3 AD-DEM, 3 CON) shows that CSNK1A1 signal is consistently observed within tau-positive neurons in AD cortex and strongly co-localises with tau pathology, while minimal neuronal CSNK1A1 is observed in controls (**Figure 7G, Supplementary Figure S6**). Our observed co-localisation is consistent with a prior study(102) and recent analyses identifying tau-CSNK1A1 proteomic correlation(103,104).

SCAI shows expression in a proportion of cells harbouring tau tangles, but is not exclusively enriched within these neurons. Instead, SCAI labels a sparse population of intensely immunoreactive subcortical cell bodies, accompanied by prominent beaded processes that extend throughout the cortex (**Figure 7F, Supplementary Figure S6**). This staining is reminiscent of interstitial white-matter neurons(105). Combined with clear membrane-association in cortical i^3^Neurons, we see a pattern that is inconsistent with the previously defined localisation and nuclear role of SCAI in cancer. Together, these observations suggest that SCAI adopts a broader subcellular distribution in human neurons than previously recognised, although its functional role in the brain and AD remains to be established.

## Discussion

Population-scale proteomics has progressively resolved molecular disease phenotypes at increasing resolution, from bulk tissue(14,106–109) to single-cell(110), to spatially resolved measurements within tissue architecture(111,112). Subcellular organisation represents a further, largely unexplored, axis of variation. Here, we show that this axis can be measured as a phenotype at cohort scale, uncovering disease biology that is not accessible through abundance measurements alone. This establishes intracellular localisation as a complementary dimension of population-scale molecular phenotyping that has its own inter-individual structure, its own disease associations, and a new capacity for discovery.

Our results indicate that disease-associated protein redistribution is not restricted to isolated localisation events, but instead converges on molecular systems whose function depends on tightly regulated spatial organisation. This is particularly evident for endolysosomal trafficking, axonal transport and RNA processing, processes that have emerged as central components of AD pathogenesis. Redistribution of adaptor proteins, the cargo-selective retromer complex and associated vesicle-sorting machinery is consistent with growing evidence that defective endosomal recycling is an early pathogenic event contributing to impaired cargo sorting and lysosomal homeostasis(113,114). Components of the AP-3 adaptor complex are among our strongest hits. Although recently identified as interactors of tau phosphorylated at T217, their role remains largely unexplored in neurodegeneration(82,115).

Endosomal trafficking is closely coupled to long-range transport, providing a potential link to the coordinated redistribution observed across dynactin and kinesin-associated proteins. Efficient trafficking of endosomes, lysosomes and APP-containing vesicles relies on tightly regulated microtubule-dependent transport, disrupted by pathological AD changes(116–120). Similarly, coordinated redistribution of multiple components of the U1-spliceosome is consistent with evidence that early spliceosomal dysfunction precedes overt amyloid and tau pathology, contributes to neuronal dysfunction, and associates with the insoluble tau proteome during disease (104,121–123). Finally, the identification of multiple components of the phosphoinositide (PI) pathway exposes its spatial organisation as a potentially underexplored feature of AD. As a central regulator of membrane identity and intracellular signalling, PI signalling integrates many of the trafficking, cytoskeletal and proteostatic processes identified here (**Figure 4**)(124).

Notably, our findings extend beyond contextualising established AD biology. By interrogating intracellular organisation orthogonally, we identify candidates that remain largely invisible to even the largest cohort-scale proteomic studies(14,17,104). SCAI exemplifies this discovery potential. Although the mechanisms of SCAI in brain remain to be determined, our follow-up analyses support a role in human brain that has not previously been characterised. We anticipate that parallel applications will facilitate discovery across a broad range of tissues and biological systems where intracellular organisation is altered.

Several considerations should be accounted for when interpreting these findings. Protein localisation is inherently dynamic, with moonlighting proteins occupying multiple compartments or exhibiting context-dependent localisation. This approach cannot distinguish whether multi-localisation occurs within or between cell types(125), which can exhibit altered morphology and proportions across disease(126). The use of curated markers is also constrained by a lack of ground truth for canonical localisation in brain – as exemplified in the case of SCAI. Our localisation prediction also reflects high inter-individual variability. Therefore, few proteins can be confidently assigned to a single subcellular compartment. Nonetheless, our out-of-sample localisation prediction recovers proteins such as SMOC1 and MDK, which are among the strongest abundance-associated markers reported across multiple large-scale AD proteomic studies (14,17,104), providing confidence that biologically relevant signals are retained irrespective of incomplete compartment resolution or multi-localised proteins. Furthermore, uncertainty in compartment assignment - whether arising from genuine multi-localisation or due to limited resolution at seven biochemical fractions - does not preclude the detection of disease-associated differential fractionation within our interaction model.

Fundamentally, fractionation cannot resolve cell type-specific localisation within heterogeneous tissue. Likewise, biochemical separation into seven fractions cannot fully distinguish closely related intracellular compartments, such as the secretory pathway, which has the most diverse proteomic signature across cell types (25). These challenges are further amplified in AD brain tissue by postmortem delay, freeze-thaw effects, and disease heterogeneity. While cell-based subcellular proteomics would provide greater resolution, current *in vitro* models incompletely recapitulate adult human neurodegeneration, particularly with respect to tau isoform expression and aggregation(127). Furthermore, inference of proteoform localisation relies on detection of peptides unique to a given molecular species, such as the MAPT R2 domain. Consequently, inference of proteoform-specific localisation is inherently constrained by peptide observability in bottom-up proteomics.

These challenges provide a rationale for complementary multi-omic integration, as demonstrated by recent methods such as LoRNA(128). Pairing comparative subcellular proteomics with single-cell transcriptomics, long-read sequencing, and phospho-omics may improve cell-type and proteoform-specific localisation inference(129,130), particularly as these data types become increasingly available deeply phenotyped cohorts such as ROSMAP(51,131). More broadly, comparative subcellular proteomics offers a complementary dimension for multi-omic integration as human atlases diversify across a range of tissues.

## Conclusion

In summary, we show that intracellular organisation is a measurable, individually varying molecular phenotype in human tissue. This trait can be modelled as a function of disease status, pathological burden, and resilience. Protein localisation adds molecular context to conventional measures of proteomic abundance, revealing disease-associated variation that remains inaccessible even to large-scale abundance studies. As population cohorts increasingly combine proteomic, genomic, and clinical data, we anticipate that subcellular localisation could become a standard axis of this integration.

## Methods

### Experiment and data acquisition

Fractionation methods were adapted from Itzhak *et al.*(22), and Kandigian *et al.*(18). The full protocol is detailed in **Supplementary SOP001**. Briefly, 200mg of dorsolateral prefrontal cortex (dlPFC) tissue per subject was homogenised by 20 strokes of a glass-Teflon homogeniser rotating at 800 rpm, 4°C. Homogenates were subjected to sequential ultracentrifugation (**Figure 1C**), comprising stepwise spins of 10 min at 1,000 ×g (P1), 10 min at 3,000 ×g (P2), 15 min at 5,400 ×g (P3), 20 min at 12,200 ×g (P4), 20 min at 24,000 ×g (P5), and 30 min at 78,400 ×g (P6), followed by collection of the final supernatant fraction (S7). This yielded seven subcellular fractions per subject.

Fractions were resuspended in PreOmics Lyse buffer for tryptic digestion (500 µL for P1, 150 µL for P2, 100 µL for P3–P6; the remaining supernatant (S7) was suspended 1:1 in PreOmics 2x Lyse buffer). Pellets were solubilised by sonication. Tryptic digestion and clean-up were performed using the PreOmics iST kit according to the manufacturer’s protocol. Dried peptides were resuspended in 50 µL 0.1% formic acid and quantified using the Pierce Quantitative Fluorometric Peptide Assay according to manufacturer’s instructions. Samples were normalised to 200ng in 20 µL and loaded onto EvoSep tips according to manufacturer’s instructions.

### LC-MS^2^ analysis

Peptides were analysed on a Vanquish Neo UHPLC System (trap and elute mode) coupled to an Orbitrap Astral mass spectrometer (Thermo Fisher Scientific; TFS) at the University of Dundee. Peptides were loaded onto a PepMap Neo Trap Cartridge (TFS, #174500), then separated using a C18 EASY-Spray HPLC Column (TFS, #ES906) using an 11.8 min elution gradient of 1–55% Buffer B (Buffer A: 0.1% FA:H_2_O; Buffer B: 0.08% FA in 80% ACN:H_2_O): 0.7min at 1–4% B, 0.3min at 4–8% B, 6.7min at 8–22.5% B, 3.7min at 22.5–35% B (1.8 μL/min), and 0.4min at 35–55% B (2.5μL/min).

Data were acquired in data-independent acquisition (DIA) mode, with both MS1 survey scans and MS2 scans collected with a 3ms maximum ion injection. Total cycle time per DIA loop was 0.6s.

MS1: Precursor spectra were recorded in profile mode at a resolution of 240,000, full scan range of 380–980m/z, AGC target of 5e6.

MS2: Precursor ions were HCD fragmented at 25% NCE, AGC target of 5e4. Fragment spectra were acquired in centroid mode across 149 non-overlapping 4 Th quadrupole isolation windows spanning 380–980m/z, full scan range of 150–2000m/z. R-F lens voltage was 40% throughout, polarity was set to positive, and source fragmentation was disabled. Full instrument settings are shown in **Supplementary S10.**

### Peptide quantification: DIA-NN processing of peptide spectra

Raw mass-spectra (TFS.raw) were converted to mzML using proteowizard msconvert(132), applying vendor peak picking settings (“peakPicking vendor snr = 1.0 peakSpace = 0.1 msLevel = 1-”). Peptide identification, filtering and quantification were performed using DIA-NN v2.0(133) **(Figure 1F)**, searching against a spectral library generated by in-silico digestion of two reference fasta files: UniProtKB/Swiss-Prot human proteome(60) (03/2025, 20421 proteins) and CCP-cRAP contaminants (02/2025, 125 proteins) acquired using camprotR(134). Sequences corresponding to MAPT (P10636) were manually removed from the CCP-cRAP reference, as tau represents a physiologically relevant protein.

Peptides were searched in match-between-runs mode using non-heuristic protein inference and UniProt identifiers to flag proteotypicity. The following default DIA-NN processing parameters were used:

- peptide length 7–30, m/z range 300–1800, and charge range 1–4, default precursor mass tolerance (optimised to ≈ ±4.0ppm)
- Fragment mass tolerance default (optimised to ± 10 ppm), m/z range 200–1800
- Modifications: N-terminal methionine excision (variable), Cysteine carbamidomethylation (fixed)
- Retention time alignment (non-linear regression algorithm) and chromatographic peak shape scoring
- Precursor and protein-level quantification (QuantUMS algorithm(135)) were filtered at Q-value ≤ 0.01 to achieve 1% FDR.

### QC and Data analysis

DIA-NN precursor and protein level quantitative matrices (FDR 1%) were used for downstream analyses, using R v4.4.3(136) in RStudio(137). Core tidyverse(138) packages were used for data wrangling, QC, modelling, and visualisation in ggplot2(139).

Data were stored as SummarizedExperiment(140) and MultiAssayExperiment(141) S4 classes.

#### Run-level QC

A single run with ≥ 80% missingness was excluded. Technical pools were removed prior to transformation and normalisation. Fractionation stability was demonstrated across subjects by principal component analysis (PCA), showing that run-wise log_2_-intensity profiles cluster by fraction. Fraction 7 (enriched for cytosolic and soluble proteins) is the most distinct cluster along PC1, likely due to increased data missingness (**Supplementary Figure S1**).

#### Precursor and protein QC

Contaminants(134) and non-proteotypic entries were removed. Raw abundance values were either log2-scaled to ensure normality throughout modelling, or sum-normalised within-subject to represent proportional fractionation shifts, independent of inter-individual abundance differences. For ML, PCA, and visualisations, all protein entries with < 0.7 global missingness (considered robustly detected) were mechanism-aware imputed (MAI) to preserve local covariance structure and limit distributional skew. This method considers missingness at random (MAR) or not at random (MNAR). The latter is a biologically informative feature and may represent low (or absence of) expression in some fractions due to protein localisation. The MAI package(142) predicts missingness as either stochastic (MAR) or intensity-dependent (MNAR) using a random forest classifier and employs a hybrid imputation strategy to the respective class. The settings MNAR_algorithm = “single”, MCAR_algorithm = “BPCA”, forest_list_args = list(ntree = 300, proximity = FALSE) were used.

### Localisation modelling

Proteins in the unimputed dataset were retained for modelling conditional on: (i) global missingness < 0.7, (ii) summed raw intensity > 500 across fractions for every subject, and (iii) ≥ 5 non-missing observations within each condition–fraction combination, in order to limit risk of bias due to non-random missingness.

The imputed dataset was also tested in parallel using the same protein set. All continuous variables were z-scaled prior to modelling to ensure model stability.

#### Variance decomposition

Per-protein variance was decomposed using linear mixed-effects models(143) of log2 - scaled intensities, using the fixed predictors Fraction, Condition, age at death, PMI, and sex as fixed effects, and a random intercept for subject. Variance explained per term was estimated by dropping each out of the full model and recording change in marginal R^2^.

#### Interaction model

The following approach was used to infer differential protein localisation by condition. All continuous variables were scaled prior to modelling.

For each protein *i* (*i = 1,…,4794*), log2-intensity was modelled as:

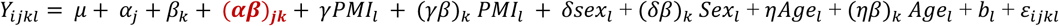

Model variables are described in **Table 3**.

**Table 3.** Mixed linear model terms.

| Term | Description |
| --- | --- |
| $Y_{ijkl}$ | Log<sub>2</sub>-intensity for protein $i$, condition $j$, fraction $k$, subject $l$. |
| $\mu$ | Model intercept; mean log<sub>2</sub>-intensity. |
| $\alpha_j$ | Fixed effect of condition: CON, RES, or AD-DEM* |
| $\beta_k$ | Fixed effect of subcellular fraction ($k=1, \dots, 7$) |
| $(\alpha\beta)_{jk}$ | Condition × Fraction interaction; term of interest used to infer condition-associated differential localisation. |
| $\gamma, PMI_l$ | Fixed effect of postmortem interval (PMI). |
| $(\gamma\beta)_{k, PMI_l}$ | PMI × Fraction interaction. |
| $\delta, Sex_l$ | Fixed effect of sex. |
| $(\delta\beta)_{k, Sex_l}$ | Sex × Fraction interaction. |
| $\eta, Age_l$ | Fixed effect of age at death. |
| $(\eta\beta)_{k, Age_l}$ | Age × Fraction interaction. |
| $b_l \sim N(0, \sigma_{2b})$ | Subject-level random intercept. |
| $\varepsilon_{ijkl}$ | Residual error. |

*\*For continuous analyses (performed on imputed data), the condition term was replaced with pathology or ‘resilience’ variables.*

Differential localisation was inferred using the Condition × Fraction interaction, which captures variation in the shape of fractionation profiles across conditions, independent of overall protein abundance.

1. *log_2_(intensity) ∼ Condition + Fraction + covariates*
2. *log_2_(intensity) ∼ Condition + Fraction + Condition:Fraction + covariates*

Two nested models: reduced (1) and full (2), were compared by likelihood-ratio test(144), whereby the resulting *X^2^*, and probability *p* < 0.05 determine whether the full model shows significantly better fit. Effect sizes were quantified as the change in marginal R^2^ (*ΔR^2^*). The Benjamini-Hochberg(72) method was used to control for false discovery rate (FDR), with an adjusted significance threshold of *p* < 0.05.

#### Longitudinal resilience model

Longitudinal global cognition was modelled using linear mixed-effects regression(143) to estimate individual rates of cognitive decline. Cognition (across study follow-up)(131) was modelled as a function of follow-up year (where year 0 represents baseline visit), adjusted for covariates: baseline global cognition score, baseline age (baseline rather than terminal measures were used to avoid conditioning on downstream consequences of the cognitive trajectories being estimated), sex, and global neuropathological burden (postmortem, used as an estimate of accumulated disease burden across the cognitive trajectory). To account for the influence of these covariates on rate of decline, each was also included in interaction with follow-up year. Subject-specific random intercepts and random slopes were included to account for within-subject correlation, capturing variation in both baseline cognition and rate of decline:

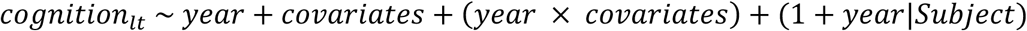

Continuous covariates were standardised before modelling and the model was fitted by restricted maximum likelihood. Subject-specific random slopes were extracted as the resilience measure. For each individual, this resilience slope represents their deviation from the trajectory predicted by the fixed covariates. For example, positive slopes represent slower than expected decline (resilience), while negative slopes represent accelerated decline (vulnerability).

Resilience-associated localisation shifts were tested by replacing diagnostic condition with these resilience slopes. Sex was not reintroduced into the localisation model. Proteomics data were collected at the age of death, so PMI and age at death were retained in the localisation model. For each protein, nested mixed-effects models with and without the resilience slope x Fraction interaction were compared by likelihood ratio test as described in the above section.

### Ontological enrichment analysis

Fractionation profile cluster Gene Ontology: Cellular Component (GO:CC)(65,66) enrichment, and enrichment of significant results were assessed using clusterProfiler(145), using a background set of all UniProt(60) entries retained after QC. Network enrichment was evaluated in the STRING(69) database.

### Ranked enrichment analysis: validation of localisation prediction

Proteins were ranked independently for each predicted compartment according to their mean out-of-sample (OOS) localisation probability across all subjects. Gene Set Enrichment Analysis (FGSEA v1.34.2)(94) was then performed on each ranked list using Gene Ontology Cellular Component (GO:CC)(65,66) annotations obtained from msigdbr v25.1.1(146). Gene sets were restricted to pathways containing between 100 and 300 detected proteins after intersecting with the background proteome. Enrichment significance was estimated using 10,000 permutations (scoreType = “pos”). Significant positively enriched pathways (FDR < 0.05) were retained for visualisation.

### Marker curation

Protein localisation markers were curated by integrating annotations from six external reference datasets; Human Cell Atlas(147), DOM(22), hyperLOPIT(148), LOPIT-DC(20), Kandigian et al. (2022)(18), and neuronal annotations from SynGO release v1.3(149) (**Figure 2C**). Source datasets were merged by UniProt(60) identifier or gene symbol, and filtered for entries present in the experimental dataset of robustly detected proteins (missingness <0.7).

A set of predefined keyword-based regular expression patterns was used to map source labels to broad compartment classes: Nucleus, Cytosol, Mitochondria, Peroxisome, Plasma Membrane, Postsynapse, Presynapse, Ribosome, Cytoskeleton, Proteasome, and Secretory pathway (Golgi, ER, endolysosome). Consensus assignments were defined as those with matching class compartment labels across ≥2 sources, while ambiguous markers (with conflicting classes) were excluded. For underrepresented classes (Postsynapse, Presynapse, Peroxisome, n < 30), unambiguous annotations with external validity (in SynGO or UniProt) were retained if consistent with their cluster GO:CC enrichment term(65,66) label from hierarchical clustering (Ward’s D2, k = 15)(64,67) of the experimental data.

### ML-based subcellular map generation

For each subject-fraction matrix of robustly detected proteins, a T-augmented Gaussian Mixture (TAGM) model(38,39) was trained (expectation-maximisation convergence, n = 100 iterations) on 782 markers representing *k* = 11 compartment classes, using sum-normalised protein fractionation profiles as features (MAI imputed data). For each *k*, MAP (Maximum-a-Posteriori) parameters of a multivariate normal distribution *N*(*μ_k_*, *Σ_k_*) were estimated: multivariate mean profile (*μ_k_*), shrinkage-regularised covariance (Σ*_k_*), mixture weights (*π_k_*), and ‘outlier probability’ multivariate t-distributed parameters. These were used to predict class membership *P*(*k* | *x*) and outlier *P*(*outlier* | *x*) probabilities of unlabelled proteins. An assignment threshold *P*(*k* | *x*) × [1– *P*(*outlier* | *x*)] ≥ 0.95 was set.

The resulting 7,264 proteins x 11 compartments matrices (whereby each matrix element represents posterior probability *P*(*k* | *x*) of a protein j in compartment k) were concatenated for *i* = 74 subjects. Subject G2 (CON) with an incomplete fractionation profile was excluded.

### Out-of-sample localisation prediction

To model subcellular localisation at cohort scale, a weighted multinomial regression framework was applied with subject-level cross-validation to account for inter-individual variability and uncertainty in TAGM-MAP predictions.

Within each disease group, subjects were partitioned into folds of five (n = 5/25). One split in CON contained 4 due to the removal of G2. A multinomial model(150) (GLM, maxit = 200) was used to predict subcellular compartment assignment (the response variable, TAGM-MAP compartment allocation) using sum-normalised fractionation profiles (proportional abundance across fractions 1–7) and subject-level covariates (age, sex, postmortem interval). For each fold, the model was trained on the remaining 20 subjects and used to generate out-of-sample predictions for those held out, ensuring that localisation probabilities for each individual were estimated independently of the data used in model training. Subject G2 (incomplete fractionation profile) did not contribute to training.

Observations were weighted by the posterior probability of the assigned compartment, such that proteins with higher localisation confidence contributed more to model fit. Curated marker proteins were not included in the model. Predicted out-of-sample probabilities were subsequently transformed to log-odds for downstream analysis of localisation distributions across disease groups. For each protein, out-of-sample localisation probabilities were averaged across subjects within each disease group to generate an 11-dimensional localisation probability distribution spanning all compartments. Jensen-Shannon divergence(151) was then calculated between disease groups to quantify differences in the overall localisation probability distribution.

### Tissue enrichment strategies

#### Cytosolic, nuclear, and synaptic

To independently validate localisation, frozen dlPFC from a subset of the ROSMAP cohort (n = 3 per condition) was subjected to sequential subcellular fractionation using an optimised protocol combining the Syn-PER Synaptic Protein Extraction Reagent and NE-PER Nuclear and Cytoplasmic Extraction Reagents (Thermo Fisher Scientific, TFS) (**Supplementary SOP002**). Briefly, tissue was Dounce homogenised on ice in Syn-PER^TM^ reagent (catalogue #87793), followed by differential centrifugation to isolate cytosolic and synaptosome-enriched fractions. The nuclear pellet generated during the initial low-speed centrifugation was subsequently processed using the NE-PER^TM^ protocol (catalogue #78833) to obtain a nuclear protein fraction. All procedures were performed at 4°C in the presence of protease inhibitors. Fractions were separated by SDS-PAGE on Bio-Rad stain-free gels (#4568096), transferred to PVDF low fluorescence membranes (Bio-Rad 162-0261), and immunoblotted with antibodies against SCAI (rabbit monoclonal antibody EPR4128, Abcam, ab124688) or CSNK1A1 (rabbit monoclonal antibody EPR1961(2), Abcam, ab108296). Bands were visualised by enhanced chemiluminescence (Bio-Rad Image Lab software) and normalised to stain-free total protein.

#### Separation of sarkosyl-insoluble material

Sarkosyl-insoluble protein fractions were isolated from a subset of ROSMAP dlPFC samples (n = 2 per condition) using an optimised sequential sarkosyl extraction and serial ultracentrifugation protocol, adapted from Arseni et al. (2024)(152). The final detergent-insoluble pellet was resuspended by sonication and analysed by immunoblotting using the same antibodies. Full experimental details are provided in **Supplementary SOP003**.

### Cortical I^3^Neurons

#### iPSC differentiation protocol

Human CRISPRi-i3N induced pluripotent stem cells (G3dCas9, NIH, USA) were differentiated into excitatory cortical neurons using an inducible NGN2 protocol (**Supplementary SOP004**) adapted from Fernandopulle et al. (2018) and Tian et al. (2019) (153,154).

iPSCs were maintained on Matrigel (Scientific Laboratory Supplies, 354277) in mTeSR1 plus medium (100-0276) before induction with doxycycline (2 μg/mL) in DMEM/F12 (21331020), non-essential amino acids (11140035) and GlutaMAX (35050061). On day 3, induced neurons (i3Neurons) were replated onto poly-L-ornithine (P3655-10MG) /laminin-coated (L2020-1mg) culture vessels and matured in BrainPhys medium (05790) supplemented with B27 (17504001), BDNF (10 ng/mL) (450-02-1MG), NT-3 (10 ng/mL) (Qk058-0100), laminin (1 μg/mL) (L2020-1MG) and doxycycline (D9891-1G). Cultures were maintained by twice-weekly half-medium changes until experimental analysis.

#### Immunocytochemistry

For characterisation of protein localisation in cortical i^3^Neurons, cells were fixed in 4% paraformaldehyde (PFA) for 10 min at room temperature (RT), washed three times with PBS, and permeabilised with 0.1% saponin in PBS for 10 minutes. Non-specific binding was blocked for 1 hour at RT using 10% normal donkey serum (430282) in PBS containing 0.1% saponin. Primary antibodies (anti-SCAI EPR4128, CSNK1A1 EPR1961(2), and anti-MAP2, ab92434-100ul chicken) were diluted in blocking solution and incubated overnight at 4°C. Following PBS washes, fluorophore-conjugated secondary antibodies (Donkey anti-rabbit Alexa 647, A31573, Alexa 555 Donkey anti-chicken, A78949) diluted at 1:1000 in PBS were applied for 1 hour at RT in the dark. Cells were washed with PBS, stained with DAPI (D9542-10MG) (1:1000 in PBS) for 10 minutes at RT, and washed again before fluorescence imaging. For G3dCas9 iPSC-derived cortical neurons, washing steps were minimised to reduce cell detachment.

### Tissue immunohistochemistry

#### Samples

Human brain frontal cortex FFPE tissue was acquired from New York University Alzheimer’s Disease Center (USA), which provides human brain tissue from an ethically approved longitudinally assessed regional brain donor program on neurodegenerative disease. Brain tissue was acquired under protocols with Institutional Review Board (IRB) approval at NYU Grossman School of Medicine. In all cases, written informed consent for research was obtained from the patient or legal guardian, and the material used had appropriate ethical approval for use in this project. All patients’ data and samples were coded and handled according to NIH guidelines to protect patients’ identities.

Sections from n = 3 clinical AD (AD-DEM) and n = 3 cognitively unimpaired age-matched controls (CON) were used for immunohistochemistry. Case-specific details are summarised in **Table 4**.

**Table 4.** Sample information for immunohistochemistry.

| Condition | Region | Age | Sex | Amyloid Semi-Quant (0–5) | AT8 Semi-Quant (0–5) |
| --- | --- | --- | --- | --- | --- |
| CTL1 | Superior Frontal Gyrus | 48 | M | 0 | 0 |
| CTL2 | Superior Frontal Gyrus | 64 | F | 0 | 0 |
| CTL3 | Superior Frontal Gyrus | 95+ | F | 0 | 0 |
| AD1 | Superior Frontal Gyrus | 90 | F | 5 | 5 |
| AD2 | Superior Frontal Gyrus | 66 | F | 4 | 5 |
| AD3 | Superior Frontal Gyrus | 85 | F | 4 | 5 |

#### Immunohistochemistry

FFPE tissue sections underwent fluorescent immunohistochemistry using the method described in (79). Briefly, sections were deparaffinised and rehydrated through a series of xylene and ethanol washes. Antigen retrieval was achieved by boiling in citrate buffer for 21 min (0.05 mM sodium citrate, 0.05% Tween-20, pH 6). Sections were blocked in 10% normal horse serum and incubated with anti-Ab (BioLegend, 4G8, 800701, 1:750), anti-Tau (ThermoFisher, PA5-95648, 1:4000), and either anti-SCAI (Abcam, ab124688, 1:500) or anti-CSNK1a1 (Abcam, ab206652, 1:100) in 4% normal horse serum overnight at 4°C. Sections were treated with TrueBlack (Cell Signaling Technology, 92401, 1x in 70% ethanol) for 1 min at RT to minimise autofluorescence. CF543-(Biotium, 20305, 1:4000), AlexaFluor594-(Jackson ImmunoResearch Laboratories, 703-585-155, 1:2000), Cy5-(Jackson ImmunoResearch Laboratories, 711-175-152, 1:2000) conjugated secondary antibodies and Hoechst 33342 (Sigma, B2261, 1:10000) were applied for 2 h at room temperature prior to cover-slipping with Antifade ProLong Glass (Invitrogen, P36984). Whole slide images were acquired using an Olympus VS200 Slide Scanner at 20x (NA 0.8) magnification. Empty channel 488 was captured to allow for autofluorescence subtraction. Representative 40× images (NA 1.25) were captured on a Leica Stellaris TauSTED Confocal microscope.

## Supporting information

Supplementary Figures S1-10, SOP001-4

## Data Availability

Model outputs are provided as Supplementary Data. Participant-level clinical and neuropathological data from ROSMAP are available through the Rush Alzheimer’s Disease Center and AD Knowledge Portal, subject to the applicable data-use agreements. Raw and processed mass spectrometry data will be deposited to the ProteomeXchange Consortium via the PRIDE repository(156) prior to journal publication.

## Acknowledgements

This work was funded by an Alzheimer’s Research UK Senior Research Fellowship (ARUK-SRF2022A-012), awarded to B.C.C, who is additionally supported by a BD^2^ Discovery Grant (DG240508). The research was carried out at the National Institute for Health and Care Research (NIHR) Oxford Health BRC and the Oxford Health BRC Molecular Targets theme. ROSMAP is supported by P30AG10161, P30AG72975, R01AG17917, R01AG015819, U01AG072572, and U01AG046152.

ROSMAP resources can be requested at https://www.radc.rush.edu and https://www.synapse.org. Immunohistochemical analyses performed at the New York University Alzheimer’s Disease Center by T.W. were supported by NIH/NIA grant P30AG066512 and U24NS141774. O.M.C was funded by MRC grant MR/Y010078/1 and a Todd-Bird JRF from New College, Oxford. We acknowledge the use of BioRender.com in generation and assembly of Figures 1–7.

## Author Contributions

H.A.J. performed computational processing and analysis of the proteomics data, generated figures, assisted with tissue fractionation experiments, interpreted the data, and wrote the manuscript. P.S. performed tissue fractionation and iPSC-neuron experiments. K.B. performed immunohistochemistry and imaging. A.J.S. and L.P. assisted with ultracentrifugation experiments. I.A. optimised the sarkosyl extraction protocol and assisted with sarkosyl experiments. B.G. performed mass spectrometric analysis. D.A.B. oversaw ROSMAP sample selection and cohort resources. T.W. provided neuropathological expertise and New York University Alzheimer’s Disease Center tissue samples. E.D. coordinated provision of New York University tissue resources and contributed to interpretation of the validation studies. S.L.F. and O.M.C. provided technical guidance, analytical oversight, and interpretation throughout the study. B.C.C. conceived the study, and performed the original subcellular fractionation experiments. B.C.C., O.M.C., and S.L.F. supervised the project, interpreted the data, and edited the manuscript. All authors reviewed and approved the final manuscript.

