## Supplementary Figures S1-10, SOP001-4 for "Population-scale subcellular proteomics reveals intracellular remodelling across the Alzheimer’s disease-resilience spectrum"

#### SOP001: Differential ultracentrifugation of 75 dlPFC samples

##### II) Pellet preparation for MS analysis (PreOmics Kit)

The PreOmics kit ensures reproducibility when this process is repeated across many samples.

1. **Resuspension & Lysis**
   a) Resuspend P1 in 500 μL 1× PreOmics Lyse buffer
   b) Resuspend P2 in 150 μL 1× PreOmics Lyse buffer
   c) Resuspend P3–P6 in 100 μL 1× PreOmics Lyse buffer each
   d) For each S7, add 100 μL 2× PreOmics buffer to 100 μL of sample (total 200uL, represents the cytosolic fraction, P7)
2. **Sonication**
   a) P1: 10 × 30 s
   b) P2–P6 and S7: 2 × 20 s
   c) Clean sonicator and area with 10% Chemgene.
3. **Protein Denaturation**
   a) Incubate in a heating block at 95 °C, 1000 rpm, 10 min.
   b) Briefly spin down at 300 rcf, 10 seconds at RT
   c) Add amount of lysate to tubes as below:

| Lysate amount : | 1x Lyse buffer |
|---|---|
| P1 10 uL | 40uL |
| P2 10 uL | 40 uL |
| P3 20 uL | 30 uL |
| P4 40 uL | 10 uL |
| P5 50 uL | |
| P6 50 uL | |
| S7 50 uL | |

4. **Digestion**
   d) In the tubes, add 210 uL of RESUSPEND to DIGEST. Shake (10 min, RT at 500rpm), then pipette up/down. 1 tube of DIGEST should last for 4 reactions
   e) Add 50 μL of DIGEST to tube, incubate in pre-heated heating block 37 °C, 500 rpm, 3 hours)
   f) Add 100 μL of STOP to tube (precipitation may occur), shake (1 min, RT at 500rpm)
   g) Spin sample in centrifuge (1 min, 16,000 rcf)
5. **Cleanup (Cartridge)**
   a) Use adapter to place a cartridge in a waste tube (label all tubes)
   b) Load S7 supernatant from 1d) onto cartridge (be careful not to damage cartridge bottom layer),
   bi) spin cartridge in centrifuge at 3800 rcf, 1–3 min (adjusted to ensure complete flow through).
   c) add 200 μL of WASH 1, repeat step 4c)
   d) add 200 μL WASH 2, repeat step 4c) *SP
   e) Use adapter to place cartridge in a fresh collection tube (label all tubes)
   f) add 100 μL ELUTE to cartridge, repeat step 4c) keep flow through in same collection tube and discard the cartridge
   g) repeat step 5bi) and keep flow-through in a collection tube
   h) Dry collection tube in a vacuum evaporator (45 °C, until completely dry); store at −20 °C (if <2 weeks) or −80 °C.

##### III) Peptide Quantification & EvoSep Loading

1. Resuspend peptides in 50 μL 0.1% formic acid (10uL FA : 10mL distilled H2O) Shake (RT at 500rpm , 5 mins)
2. Quantify peptide concentration (write down) with Thermo Flourescent Peptide Quant kit (according to manufacturers instructions)
3. Adjust to 200 ng in 20 μL for loading on EvoSep tips
4. Follow EvoSep manufacturer protocol for loading onto tips
5. Transport to proteomics facility for Analysis

Revision: SOP004 v1.0 | Effective: 04/09/23 | Review date: 04/09/24
Author: Becky Carlyle

#### SOP001: Preparation of Frozen Brain Tissue for Subcellular Fractionation and Proteomic Analysis

##### Summary

This protocol details the process of reproducibly homogenizing and fractionating frozen brain tissue samples (200 mg of human dorsolateral prefrontal cortex (dlPFC)) into subcellular fractions, followed by sample preparation with the PreOmics kit for proteomic analysis by mass-spectrometry.

##### Reagents for Solution Preparation

| Reagent | Stock | Amount | Final Concentration |
|---|---|---|---|
| **Sucrose powder** | | 51.345g to 50mL | 3 M *Note: do not put on ice – heat to dissolve / adjust conc. |
| **Lysis Buffer (100 mL)** | | | |
| • Tris-HCl, pH 7.5 | 1 M | 2.5 mL | 25 mM |
| • Sucrose | 3 M | 1.666 mL | 50 mM |
| • $MgCl_2$ | 1 M | 50 μL | 0.5 mM |
| • EGTA | 0.5 M | 40 μL | 0.2 mM |
| • Complete EDTA-free protease inhibitors | Tablet | 2 tabs | |
| • Water (dH2O) | | to 100 mL | |
| **2.5 M Sucrose Buffer (10 mL)** | | | |
| • Sucrose | 3 M | 8.33 mL | 2.5 M |
| • Tris-HCl, pH 7.5 | 1 M | 250 μL | 25 mM |
| • $MgCl_2$ | 1 M | 5 μL | 0.5 mM |
| • EGTA | 0.5 M | 4 μL | 0.2 mM |
| • Water (dH2O) | | to 10mL | |

##### Equipment

- Teflon-glass homogeniser
- Refrigerated centrifuge (up to 1,000 ×g)
- Ultracentrifuge with Type 70.1Ti rotor and Ultraclear Beckman tubes
- PreOmics iST sample prep kit
- Heating block with shaking capability
- Vacuum evaporator
- EvoSep tip loading system

##### I) Subcellular Fractionation

1. **Homogenisation**
   - Add 4 mL lysis buffer per 100 mg tissue/cells
   - Homogenise in glass Teflon homogeniser: 20 strokes at 800 rpm
   - This renders samples acellular. HTA Act no longer applies – record date this step was performed in sample log
2. **Nuclear Pellet (P1)**
   - Transfer 8 mL of the lysate to 15 mL-sized falcon tube
   - Add 2.5 M sucrose buffer to a final concentration of 250 mM (640 μL/8mL lysate)
   - Mix gently, centrifuge 1000 ×g, 10 min, 4 °C.
   - Allow pellet to settle. Remove as much supernatant as possible and transfer to an Ultraclear Beckman ultracentrifuge tube
   - Collect pellet 1 (P1), snap freeze at -80C; retain supernatant for subsequent steps
3. **Serial Centrifugation**

- For all steps, spin at 4 °C, allow to settle, remove supernatant and transfer to a new ultracentrifuge tube. Keep supernatant on ice while you snap freeze pellet at -80C.

| Pellet | g-force (xg) | Time (min) | 70.1Ti Rotor Speed (rpm) |
|---|---|---|---|
| P2 | 3000 | 10 | 6600 |
| P3 | 5400 | 15 | 8854 |
| P4 | 12200 | 20 | 13309 |
| P5 | 24000 | 20 | 18666 |
| P6 | 78400 | 30 | 33737 |

- The final supernatant 7 – S7, should be kept on ice.
- Store all pellets at −80 °C. Clean rotor with 70% ethanol, dry well.

### SOP002: Syn-PER/NE-PER fractionation

**SOP008 – Preparation of frozen brain tissue for sub-cellular fractionation and analysis by western blot**

Couple Syn-PER (87793) with NE-PER (78833) kits from ThermoFisher

**Cleaning and decontamination:**

- All single-use plasticware that comes into contact with samples derived from human brain tissue should be collected in a labelled Dispo-jar. When full, the Dispo-jar should be labelled with Biohazard tape, the date and group, and placed in the yellow box for disposal through the clinical waste route
- Re-usable tools such as the steel dissection plate, mallet and forceps should be submersed in 10% Chemgene for 30 minutes before rinsing with water and drying
- Wipe the homogenizers (sonication or manual) and fume hood surface with 10% Chemgene after use
- Lysates made from human tissue in the lab are all small volume below 10 mls. In case of spillage, flood the area with 10% Chemgene for 30 minutes, then wipe up.

**Step 1 – Subcellular fractionation.**

- Add protease inhibitors to the required volumes of reagents (Syn-PER, CERI, and NER). Stock: 100× (use mini-complet tablets dissolved in 200 μL PBS, or Halt Protease Inhibitor Cocktail, EDTA-free, 100×, 1 mL).
- Perform all steps -including homogenization and centrifugation- at 4 °C to minimize proteolysis, dephosphorylation, and denaturation. Keep samples and extracts on ice.
- Place the Dounce tissue grinder on ice before use.
- Pre-chill low-protein-binding tubes.
- Always keep brain samples on dry ice until just before homogenization.
- Perform homogenization in a fume hood.
- Save an aliquot of every supernatant, including wash fractions, for Western blotting.

**Protocol**

**Procedure for Synaptic Protein Extraction from Neuronal Tissue**

1. Weigh brain tissue samples. Add 10mL of Syn-PER reagent per gram of tissue (e.g., 2mL of Syn-PER Reagent per 200mg of brain tissue).
2. Perform Dounce homogenization on ice with ~10 slow strokes.
3. Transfer homogenate to an appropriate centrifuge tube(s).
4. Centrifuge the tube at 1200 × g for 10 minutes at 4°C. Keep the nuclear pellet on ice to perform nuclear proteins extraction* and transfer supernatant to a new tube (this is a mix of cytoplasmic and synaptic proteins). If required, save a sample of the supernatant (homogenate) for analysis.

These steps render the samples acellular. HTA Act no longer applies after this point. Record the date this step is performed in the sample log in the OneDrive folder

5. Centrifuge supernatant at 15,000 × g for 20 minutes at 4°C.
6. Remove the supernatant from the synaptosome pellet. If required, save the supernatant (cytosolic fraction) for analysis.
7. Add 1-2mL of Syn-PER Reagent per gram of sample to suspend the synaptosome pellet (e.g., 500μL for 200-400mg of brain tissue).

Note: Recommended volumes should result in 3-4μg/μL of synaptic protein.
8. Maintain the synaptosome suspension on ice until performing neurotransmitter release studies or downstream applications.
Note: The synaptosome suspension can be stored in 5% (v/v) DMSO at -80°C or in liquid nitrogen for extended periods of time; however, a substantial reduction in synaptosome viability occurs after prolonged storage. For best results, perform activity studies with new synaptosomes.

**Procedure for Cytosol and Nuclear Protein Extraction**

*Nuclear pellet from Step 4: treat it as brain homogenate and start NE-PER protocol from the first step

8. Weight pellet and add CERI reagent accordingly:

| Tissue Weight (mg) | CER I (μL) | CER II (μL) | NER (μL) |
|---|---|---|---|
| 20 | 200 | 11 | 100 |
| 40 | 400 | 22 | 200 |
| 80 | 800 | 44 | 400 |
| 100 | 1000 | 55 | 500 |

*Different tissue types may require more or less NE-PER Reagents per weight to optimally extract cytoplasmic and nuclear proteins.

9. Vortex the tube vigorously on the highest setting for 15 seconds to fully suspend the cell pellet. Incubate the tube on ice for 10 minutes.
10. Add ice-cold CER II to the tube.
11. Vortex the tube for 5 seconds on the highest setting. Incubate tube on ice for 1 minute.
12. Vortex the tube for 5 seconds on the highest setting. Centrifuge the tube for 5 minutes at maximum speed in a microcentrifuge (~16,000 × g).
13. Immediately transfer the supernatant (cytoplasmic extract: in this case mainly mitochondrial proteins) to a clean pre-chilled tube. Place this tube on ice until use or storage.
14. Wash pellet with ice-cold PBS and centrifuge the tube for 5 minutes at maximum speed in a microcentrifuge (~16,000 × g).
15. Suspend the insoluble (pellet) fraction produced in Step 4, which contains nuclei, in ice-cold NER.
16. Vortex on the highest setting for 15 seconds. Place the sample on ice and probe sonicate for 20 seconds twice with an interval of 20 seconds on ice.
17. Centrifuge the tube at maximum speed (~16,000 × g) in a microcentrifuge for 10 minutes.
18. Immediately transfer the supernatant (nuclear extract) fraction to a clean pre-chilled tube. Place on ice.
19. Store extracts at -80°C until use.

**Step 2 – Sample preparation and Western blot**

**Electrophoresis:**

20. Mix 15 μL of each sample with 5 μL Laemmli buffer (4×, no DTT). Adjust with Milli-Q water as needed.
21. Denature at 95 °C for 5 min; cool to room temperature (RT).
22. Briefly centrifuge to collect condensate.
23. Fill the gel tank with ~500 mL 1× running buffer.
**10× stock recipe (make 1 L):** 30.3 g Tris base, 144.1 g glycine, 10 g SDS; bring to 1 L with Milli-Q water. Dilute 1:10 for 1×.
24. Load 5 μL Precision Plus ladder in lane 1. Use a 4–20% gel (15-well).
25. Load samples (typical volumes): 13 μL per well for 10-well mini gels; ~5 μL per well for 15-well mini gels.
26. Run at ~90 V until samples fully enter the gel, then raise to ~120 V. Stop when the dye front reaches mid-to-end of the gel.

**Semi-Dry Transfer:**

27. Open the cassette and release the gel.
28. From the cold room, open a semi-dry transfer pack; place the shiny side into the transfer cassette.
29. Place the gel on top of the shiny side.
30. Place the remaining filter paper section on top of the gel.
31. Roll gently to remove bubbles.
32. Secure the lid and place the cassette in the transfer unit.
33. Run protocol "**MariLostMe**" (11 min transfer).
34. Remove the membrane from the stack.

**Visualisation of Membrane**

35. Place the membrane in a suitable container (cut if needed).
36. Block at RT for 1 h with 5% milk in PBS (e.g., 0.75 g milk in 15 mL PBS).
37. Discard block; add primary antibody diluted in 5% milk in PBS-T (e.g., 3 μL antibody in 15 mL). Incubate overnight at 4 °C on a rocker.
38. Wash 4× for 5 min each in PBS-T.
39. Add HRP-conjugated secondary antibody diluted in 5% milk in PBS-T (e.g., anti-mouse HRP or anti-rabbit HRP, 1:5000 → 3 μL in 15 mL).
40. Incubate at RT for 1 h on a rocker.
41. Wash 4× for 5 min each in PBS-T.
42. Prepare ECL substrate: mix equal parts of the two reagents (e.g., 7.5 mL + 7.5 mL = 15 mL per membrane; ~5 mL for smaller membranes).

### SOP003: Sarkosyl-extraction

**SOP: Extraction of detergent insoluble fraction from human brain**

*Round 1 performed 21st Jan 2026*
*Subject samples G4 (AD1) F4 (CON1) B4 (RES1)*
*Round 2 performed 23rd March 2026*
*Subject samples C4 (AD2) C6 (CON2) I8 (RES2)*

**Time:** 10-12 hours
Book ahead of time **[all day]:** Hood, ultracentrifuge, thermal shaker

**Equipment:**
- 15 thick-wall 1ml polycarbonate (Beckman 11x34mm, 343778) tubes (for n=3 samples [AD/RES/CON], 3x4 spins + 3 spare)
- 1.5ml and 0.5ml protein loBind tubes
- Thermal shaker (polycarbonate tubes should fit in this)
- Ultracentrifuge, 120.2Ti rotor

**Extraction Buffer (EB):**
For ~40-50mg of sample per condition [AD/RES/CON, n=3]

| Reagent | Stock | Required | For 120ml | For 20ml | Sarkosyl |
|---|---|---|---|---|---|
| Tris-HCl, pH7.4 | 1M | 10mM | 1.2ml | 200ul | |
| NaCl | 5M | 0.8M | 19.2ml | 3200ul | |
| Sucrose | - | 10% | 12g | 2g | |
| EGTA | 0.5M | 1mM | 240ul | 40ul | |
| DTT | 1M | 1mM | 120ul | 20ul | |
| Pearce protease + phosphatase inhibitor (1) | 100x | 1X | 1.2ml | 200ul | |
| Ultrapure distilled water (invitrogen) | | | Add 98ml to get up to 120 | Add 16.34ml to get up to 20ml | |
| Sarkosyl<br>*After homogenisation* | 30% | 2% | - | Allow for: 6ml used (~1600x3 samples) | **2% buffer:** 114.3ul sarkosyl for 1600ul EB (note these may differ depending on initial sample mass) |
| *Remaining steps* | 30% | 1% | | 14ml remaining | **1% buffer:** 466.7ul sarkosyl for 14ml EB |

*EB prep comment: In round 1, we made 5ml EB then added 16ml Invitrogen ultrapure distilled water, so ended up with 21ml working solution. 14ml was kept for downstream steps at 1% sark, and 6ml of this was used for initial homogenisation (sark added after homogenisation to 2%)

Additional notes:
- Ensure thermal shaker is set to 37°C and ultracentrifuge is set at 25°C
- Pre-label all tubes by colour.
- When ultracentrifuging, draw a small dot in the respective condition colour at the top side of the rim so you know which side the pellet is if it is hard to see.
- Note the rotor position for each sample in case the pen comes off in the ultracentrifuge (you can re-draw the dot on top of rim before removal if this happens).
- Each time supernatant is collected, keep a record of how much per sample before equalising.

**Protocol:**
**Homogenisation:**
1. Weigh out approximately ~40g tissue using sterile forceps in centrifuge tubes (keep samples on dry ice until homogenisation in EB to avoid thawing). The samples weighed out were:
   - Round 1) CON: 46mg, AD: 48mg, RES: 67mg
   - Round 2) CON: 65mg, AD: 51mg, RES: 44mg
2. Homogenise samples in 40 volumes of EB (e.g. 1600ul EB : 40mg tissue) using a Dounce homogeniser. Start with CON → RES → AD, sterilising the homogeniser in-between with ethanol and letting it evaporate entirely. Due to mass variation, samples were split with the following amounts of sark EB to achieve more balanced initial concentrations:
   - Round 1) CON: 1600ul EB, AD: 1640ul EB, 1680ul EB.
     - We did ~15 strokes (CON was hard to homogenise so required 20, high myelin content?)
   - Round 2) CON: 1680ul EB, AD: 1640ul EB, 1600ul EB.
3. Add 30% stock sarkosyl to both to achieve 2% final concentration (see following amounts)
   - Round 1) CON: 114.3ul sark:1600ul EB, AD: 117.2ul sark :1640ul EB, RES: 120ul sark:1680ul EB.
4. Split out each sample into 2 thick-wall polycarbonate 1ml tubes (to fit in the rotor).
   - Round 1) CON: 800ul x2, AD: 820ul x2, RES: 840ul x2.
   - Round 2) CON: 840x2, AD: 820ul x2, RES: 800ul x2.

   These were equalised to 1mL with EB prior to centrifugation.
5. Incubate for 1 hour at 37°C with orbital shaking at 700RPM

**Wash 1 (Debris removal):**
6. Centrifuge (120.2Ti rotor) at 27,000g for 10min at 25°C. This removes high density debris.
   - Refer to additional notes here: ensure coloured dot on rim and rotor positions noted.
7. Collect supernatant with care (do not disturb pellet) and add directly to 1ml tube for next step, keeping the volume equal for each tube (volume can be equalised using EB) so that they are balanced during ultracentrifugation. The following amounts of supernatant were collected in this step
   - Round 1) CON: 770ul x2, AD: 800ul x2, RES: 810ul x2
   - Round 2) CON: 770ul x2, AD: 750ul x2, RES: 700ul x2

**Spin 1 (produces soluble fraction):**
8. Ultracentrifuge (120.2Ti rotor) at 166,000g for 30min at 25°C.
9. Remove the supernatant (sarkosyl-insoluble fraction), avoiding any myelin which may have floated to the top.
   - Add supernatant to soluble labelled tubes (e.g., per sample make 2 aliquots of 250ul + 1 aliquot with the remainder for BCA) and immediately freeze on dry ice to store at -70°C.

**Wash 2 (obtain supernatant):**
10. Add 500ul of 1% sarkosyl EB to each pellet and soften them by incubating for 1 hour at 37°C with orbital shaking at 300RPM.
11. Transfer the pellets to 1.5ml loBind tubes (split samples are combined again here) – you can gently wash until you see the pellet lift, but be careful it does not stick to the pipette or you will lose it.
12. Sonicate pellets continuously (in low attachment tubes) for 5min at 50% amplitude (room temp) using a Qsonica Q700.
    - Note: We had to split the samples out here (2x500ul) into smaller LoBind tubes. We pooled the samples back together after this back into 1ml thick wall polycarbonate tubes for further centrifugation so we went from 6 tubes (2 per condition) to 3 (1 per condition)
13. Benchtop centrifuge the pellets at 17,000g for 25min at 25°C (use 3 spare 1ml tubes with equal volume of water to balance), then collect supernatants intothick-wall 1mL ultracentrifuge tubes (you won't need to split out here if supernatant volume is <1mL). Equalise samples using 1% sarkosyl EB if necessary.

**Two further serial ultracentrifugation-incubation steps are performed hereon, followed by a sonication step to obtain the sark-insoluble final pellet.**

**Spin 2:**
14. Ultracentrifuge (120.2Ti rotor) at 166,000g for 30 mins at 25°C (remember to equalise volumes).
15. Remove supernatant and add 500ul 1% sarkosyl EB to pellet.
16. Incubate pellet for 30mins at 37°C with orbital shaking at 200RPM to soften pellets.
17. Resuspend pellets and top up to 1mL with 1% sarkosyl EB (add another 500uL)

**Spin 3:**
18. Ultracentrifuge (120.2Ti rotor) at 166,000g for 30min at 25°C (remember to equalise volumes).
19. Remove supernatant carefully (do not disturb or lose the pellet!) and add 25ul PBS to the pellet (this will be the final sark-insoluble fraction).
20. Incubate for 30mins at 37°C with orbital shaking at 200RPM to soften pellets.
21. Sonicate pellets continuously (in low attachment tubes) for 5min at 50% amplitude (room temp) using a Qsonica Q700.
22. Finally, aliquot the 25ul of insoluble fraction into 5x5ul aliquots per sample and immediately freeze on dry ice and store at -70°C.

### SOP004: iPSC differentiation

**SOP: Differentiation of CRISPRi-i3N iPSCs into Cortical Neurons**

**Cell line:** G3dCas9 (CRISPRi-i3N) human iPSCs
**Based on:** Fernandopulle et al., 2018; Tian et al., 2019

#### Stage 1. Maintenance of iPSCs (Day -1 to 0)

**Materials**

- Matrigel (Corning)
- mTeSR1 medium
- ROCK inhibitor (10 μM)
- PBS ($Ca^{2+}$/$Mg^{2+}$ free)
- Accutase
- DMSO
- DMEM KO (or equivalent)

**Procedure**

1. Coat culture vessels with Matrigel and incubate at 37°C for at least 1 hour (or overnight).
2. Prepare complete mTeSR1 medium according to the manufacturer's instructions.
3. Thaw cryopreserved iPSCs rapidly in a 37°C water bath and immediately transfer cells to 10 mL DMEM KO (or PBS containing 5% medium).
4. Centrifuge cells at 300 × g for 5 minutes.
5. Aspirate the supernatant and resuspend the pellet in mTeSR1 supplemented with 10 μM ROCK inhibitor.
6. Plate cells at:
   - 1 × $10^6$ cells/well (6-well plate)
   - 3.5 × $10^6$ cells (10 cm dish or T75 flask)
7. Replace medium daily. Remove ROCK inhibitor after 24 hours once colonies have recovered.
8. Maintain cultures until approximately 80% confluency, passaging using EDTA or Accutase as required.

#### Stage 2. Neuronal Induction (Day 0-3)

**Induction medium**

DMEM/F12 supplemented with

- N2 (1×)
- MEM Non-essential Amino Acids (1×)
- GlutaMAX (1×)

Immediately before use add

- Doxycycline 2 ug/mL
- ROCK inhibitor 10 uM

**Procedure**

1. Coat culture vessels with Matrigel and incubate for at least 1 hour at 37°C.
2. Wash iPSC cultures once with PBS.
3. Dissociate cells with Accutase for 5 minutes at 37°C.
4. Neutralise Accutase with basal medium and collect cells.
5. Count viable cells.
6. Centrifuge at 300 × g for 5 minutes.
7. Resuspend cells in complete induction medium.
8. Plate cells at
   - 1.5 × $10^6$ cells/well (6-well)
   - 8–10 × $10^6$ cells/T75
   - 25 × $10^6$ cells/T175
9. Replace induction medium daily.
10. Remove ROCK inhibitor after 24 hours while maintaining doxycycline throughout the induction period.
11. Continue induction until Day 3. Neurite extension should be visible by Day 1 and well developed by Day 3.

#### Stage 3. Neuronal Maturation (Day 3 +)

**Plate coating**

Day 0

1. Coat culture plates with poly-L-ornithine (PLO).
2. Incubate overnight.

Day 1

3. Wash coated plates three times with sterile water.
4. Allow plates to air dry.

Day 2

5. Coat plates with laminin (10 ug/mL) diluted in ice-cold PBS.
6. Incubate overnight at 37°C.

**Maturation medium**

BrainPhys supplemented with

- B27 (1×, with Vitamin A)
- BDNF 10 ng/mL
- NT-3 10 ng/mL
- Laminin 1 ug/mL
- Doxycycline 2 ug/mL
- ROCK inhibitor 10 uM (first 24 h only)

**Procedure**

1. On Day 3, dissociate induced neurons with Accutase.
2. Generate a single-cell suspension by gentle trituration.
3. Count viable cells.
4. Plate neurons at the required density:

| Plate format | Cell density |
|---|---|
| 384-well | 12,000–12,500 cells/well |
| 96-well (half area) | 25,000 cells/well |
| 96-well | 50,000 cells/well |
| 24-well | 500,000 cells/well |
| 12-well | 1 × $10^6$ cells/well |
| 6-well | 2 × $10^6$ cells/well |

5. Maintain neurons in maturation medium with twice-weekly half-medium changes.
6. Remove ROCK inhibitor after 24 hours.
7. Continue culture until the desired experimental endpoint. Doxycycline may be maintained throughout differentiation if continued NGN2 expression is required.

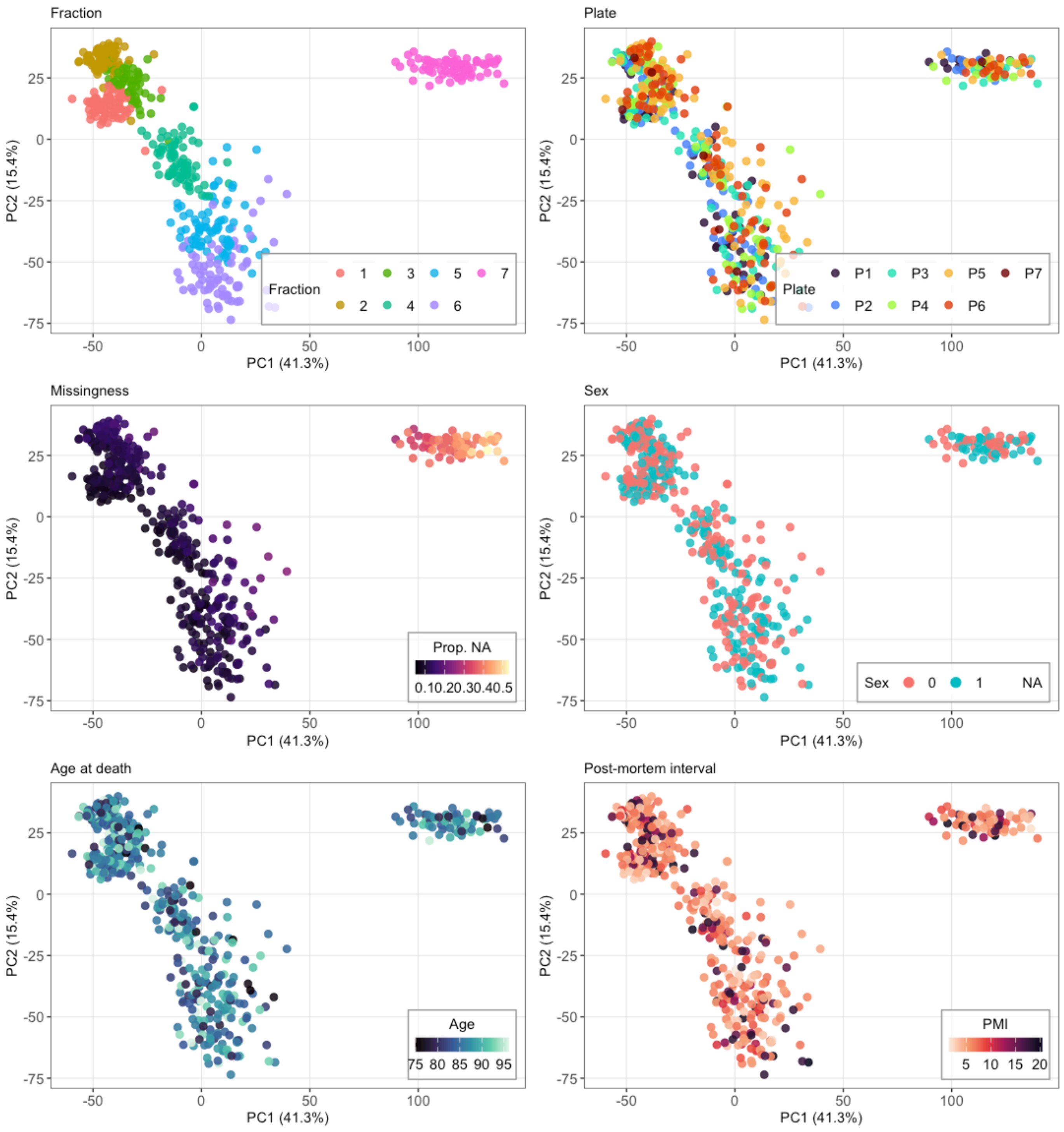


**Supplementary figure S1.** *Run-wise PCA of $log_2$-scaled protein intensities (imputed matrix) whereby samples (524/525 of LC-$MS^2$ runs passing QC) cluster by fraction, mainly along PC2. Plots are shaded by technical and cohort-level variables. Proportion of missing values in the unimputed matrix is highest in fraction 7, which shows distinct separation along PC1.*

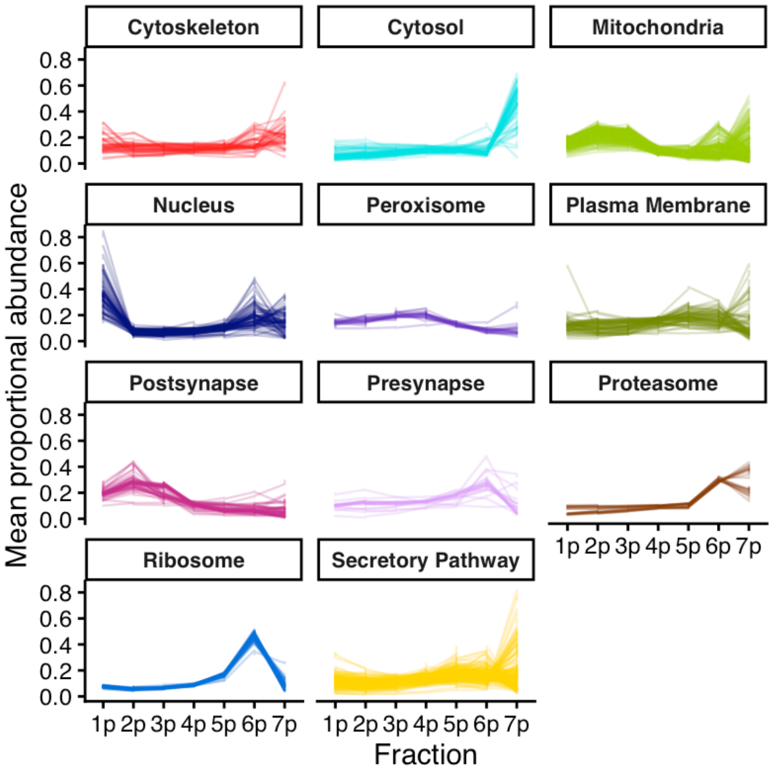


**Supplementary figure S2.** *Mean (across subjects) proportional (sum-normalised) fractionation profiles of all curated markers, grouped by subcellular compartment.*

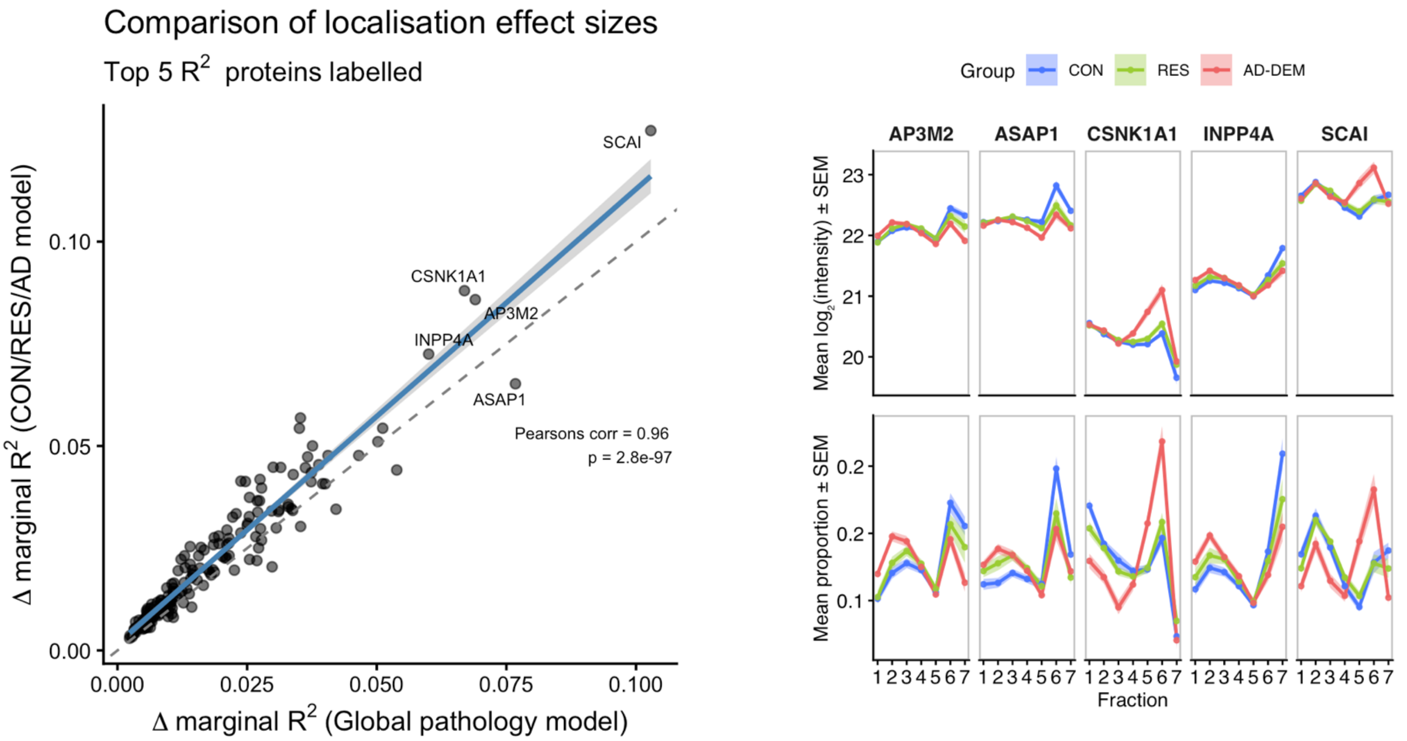


**Supplementary figure S3.** *Left: in the interaction model, the use of global pathological burden yields similar results to the CON, RES, AD-DEM categorisation. Protein effect sizes are highly correlated for the result intersection. Right: for proteins such as AP3M2, ASAP1, and INPP4A, RES has an intermediate fractionation profile between CON and AD-DEM. By contrast, CSNK1A1 and SCAI show highly divergent AD-DEM profiles, with a less distinguishable difference in CON and RES.*

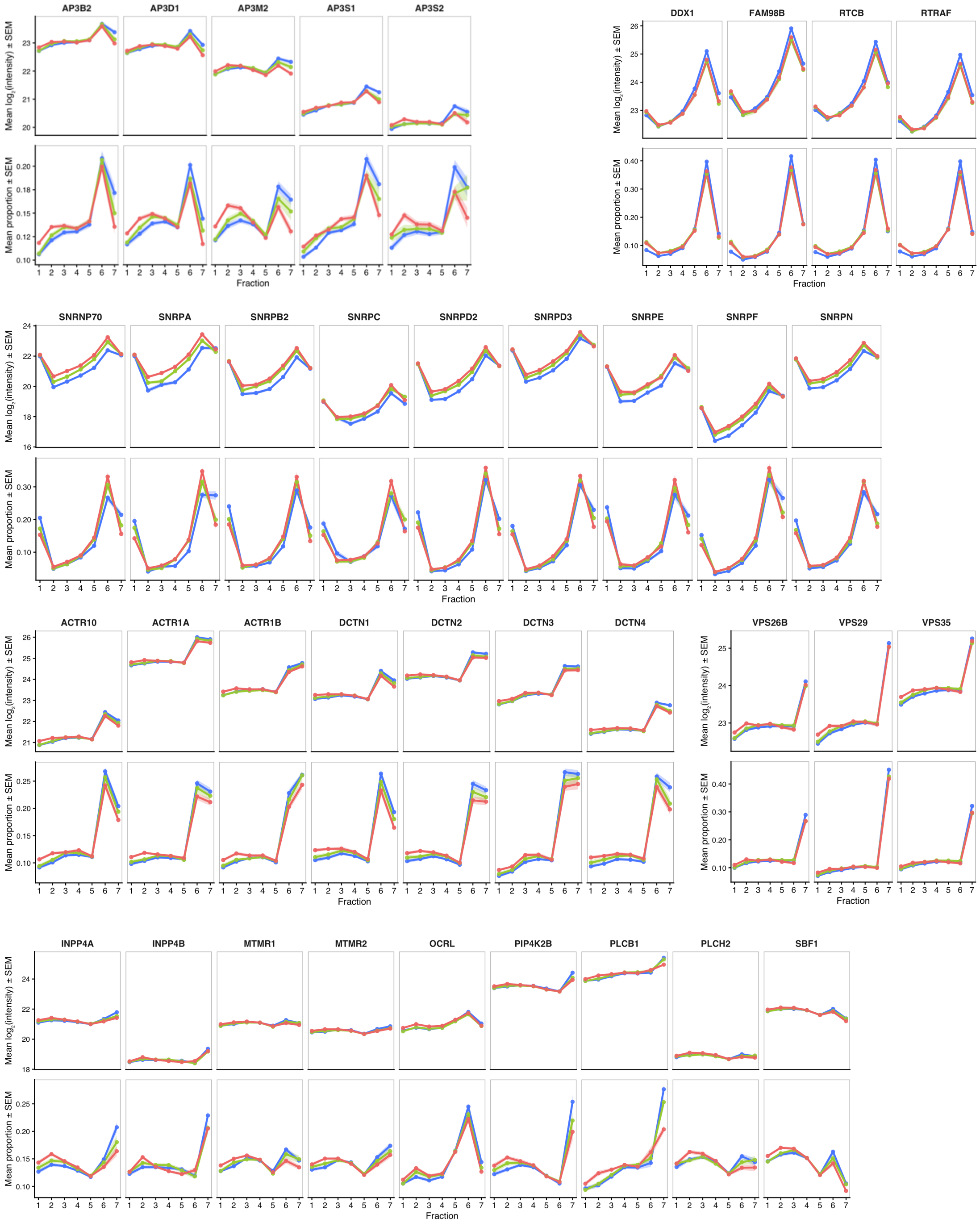


**Supplementary figure S4.** *Concordant fractionation differences observed across disease for the dynactin complex (DCTN1–4, ACTR1A/B and ACTR10), the AP-3 adaptor complex (AP3D1, AP3B2, AP3M2, AP3S1 and AP3S2), the cargo-selective retromer (VPS29, VPS35 and VPS26B), the tRNA ligase complex (DDX1, RTRAF, RTCB and FAM98B), 14-3-3 proteins (YWHAH, YWHAB and YWHAG), eight components of the U1 spliceosome, and phosphoinositide metabolism regulators (INPP4A/B, PLCB1, PLCH2, OCRL, MTMR1/2, PIP4K2B and SBF1).*

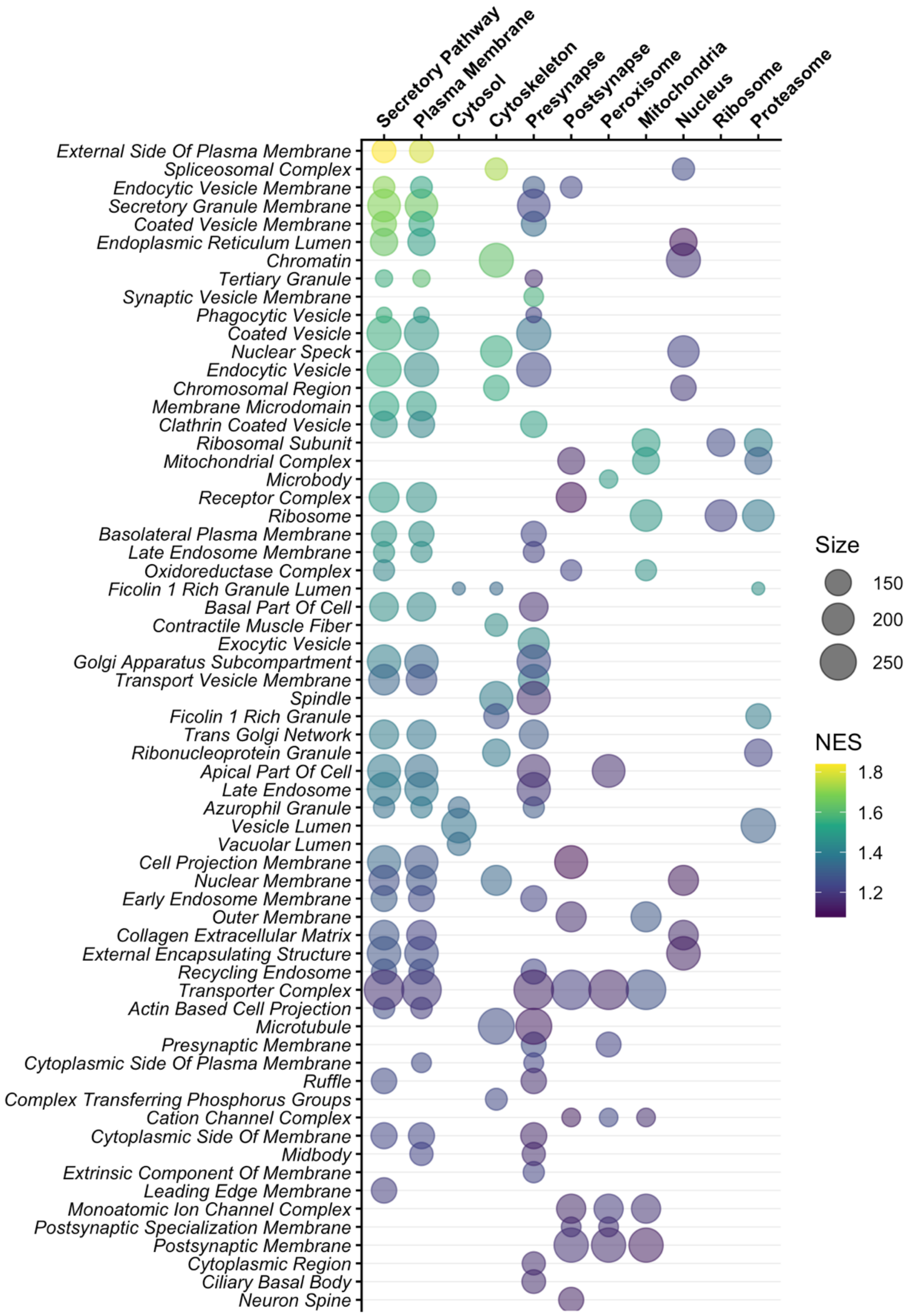


**Supplementary figure S5.** *Bubble plot of ranked set enrichment (FGSEA)(65,92,144) of proteins ordered by out-of-sample (OOS) localisation probability for each compartment (x-axis). Ranking is not conditional on an assignment probability ≥0.95, and therefore reflects whether the most likely compartment assigned by the OOS model is able to capture canonical subcellular structure. Bubble size reflects number of proteins in the GO:CC pathway, while colour reflects the normalised enrichment score (NES). Large macromolecular assemblies such as the spliceosome and ribosome are enriched across multiple compartments, possibly due to shared fractionation behaviour or compartment substructure.*

**AD2 sample:** SCAI IHC regions selected for 40X confocal imaging, highlighted below

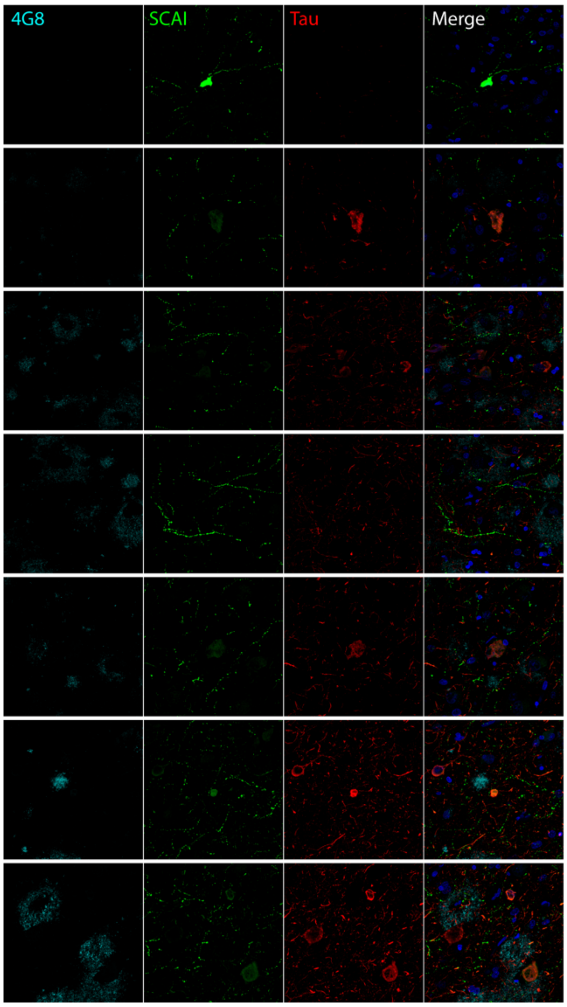


**Region 1:** Bright SCAI cell body in white matter

**Region 2:** Tangle and SCAI positive cell. SCAI intensity is low at these settings

**Region 3:** Several tangle and SCAI positive cell. SCAI intensity is low at these settings

**Region 4:** Intense SCAI beaded processes

**Region 5:** Tangle and SCAI positive cell. SCAI intensity is low at these settings

**Region 6:** Tangle and SCAI positive cell

**Region 7:** Tangle positive cells with variable levels of SCAI intensity

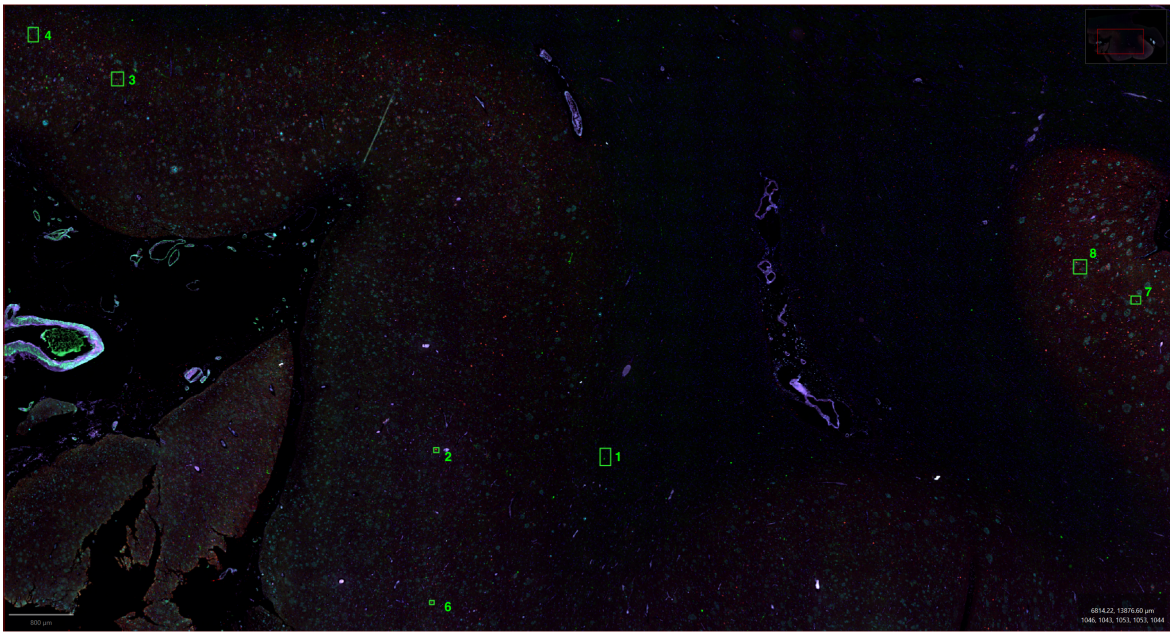

**AD2 sample:** CSNK1A1 IHC regions selected for 40X confocal imaging, highlighted below

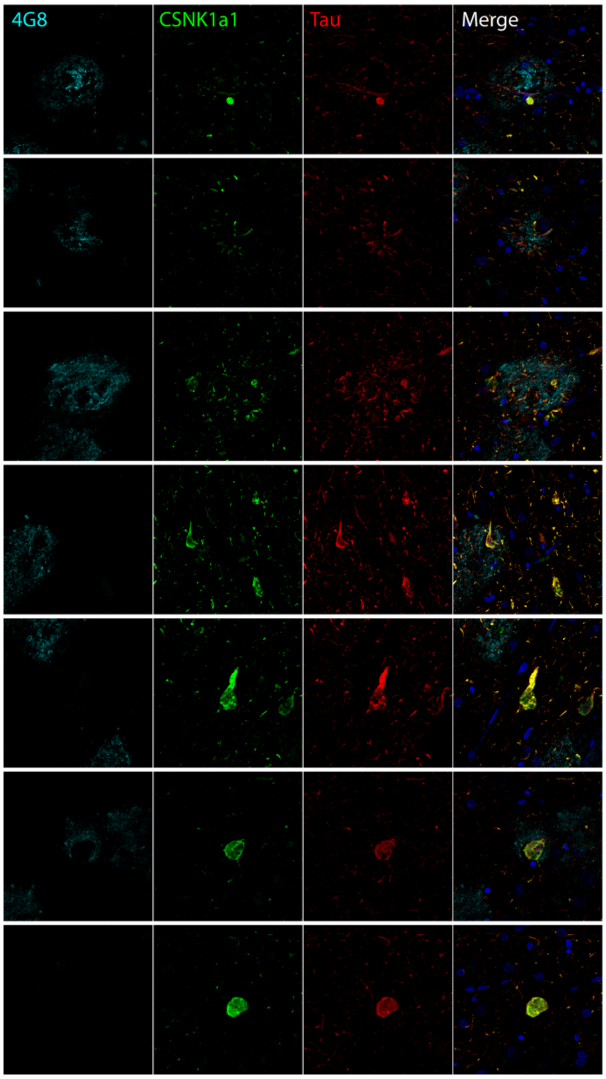


**Region 1:** Plaque with Tau/CSNK1A1 co-localisation

**Region 2:** Plaque with Tau/CSNK1A1 co-localisation

**Region 3:** Neuritic plaque

**Region 4:** Tangle positive cells with Tau/CSNK1A1 co-localisation

**Region 5:** Tangle. Image taken with a tighter z-stack (0.25um) to show texture

**Region 6:** Tangle and CSNK1A1 positive cell

**Region 7:** Tangle and CSNK1A1 positive cell

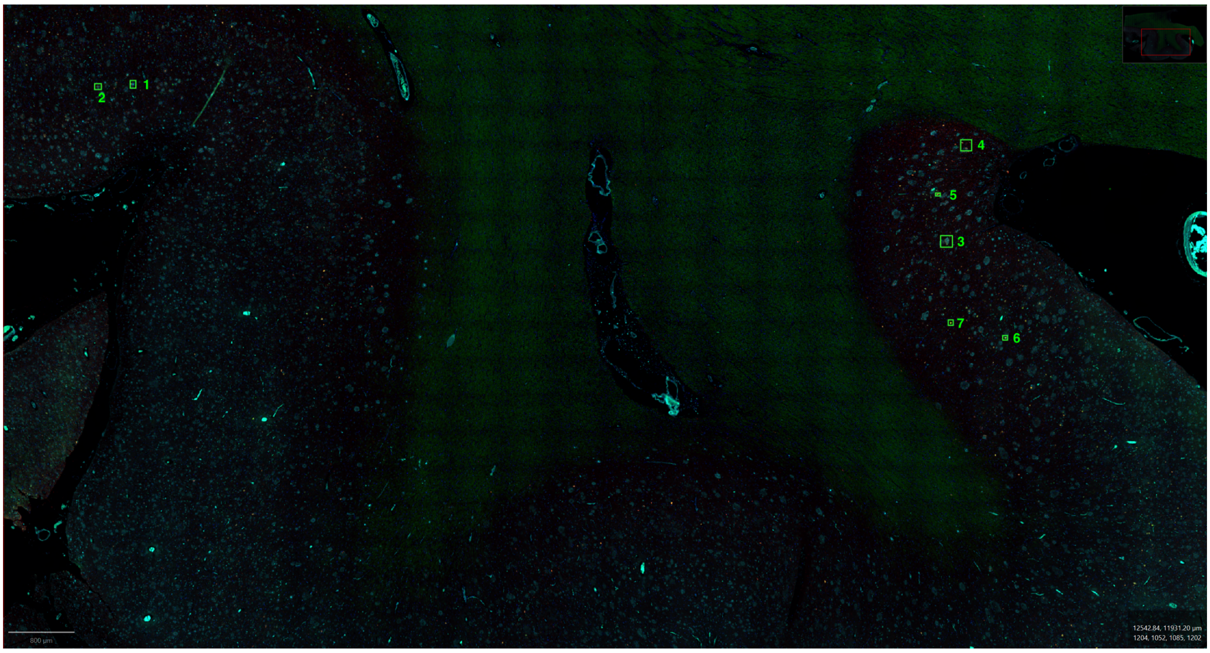

**CTL2 sample:** SCAI IHC regions selected for 40X confocal imaging, highlighted below

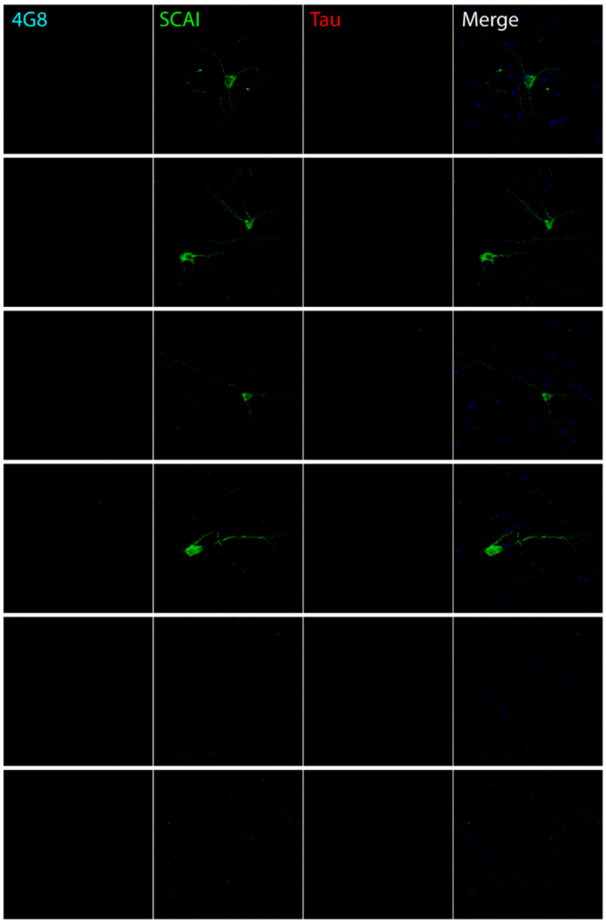


**Region 1:** SCAI positive cell body in white matter

**Region 2:** SCAI positive cells in white matter

**Region 3:** SCAI positive cell body in white matter

**Region 4:** SCAI positive cell body in grey matter

**Region 5:** SCAI processes in grey matter. Note that these are hard to see, as intensity is much lower so that bright cells (regions 1–4) are not over-saturated

**Region 6:** SCAI processes in grey matter. Note that these are hard to see, as intensity is much lower so that bright cells (regions 1–4) are not over-saturated

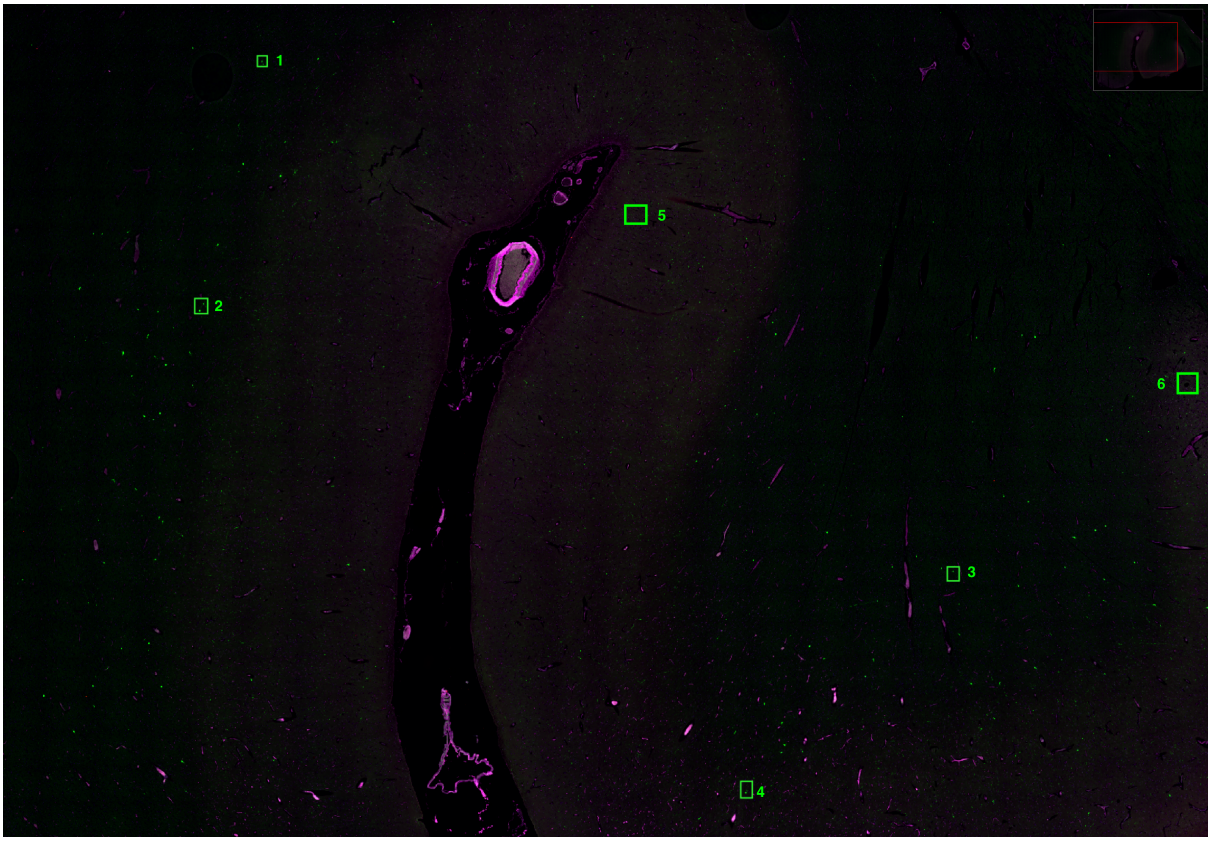

**Supplementary Figure S6. Immunohistochemical localisation of SCAI and CSNK1A1 in human cortex.** *Overview images and selected regions from multiplex immunofluorescence of AD and control cortex used for subsequent confocal imaging at 40× magnification. In AD tissue, SCAI is detected in a subset of tau-positive neuronal cell bodies but is not restricted to these cells, with prominent beaded processes also observed throughout the cortex. A sparse population of intensely SCAI-immunoreactive cell bodies and processes is also visible in white matter. CSNK1A1 signal is observed within tau-positive neurons and in regions containing neuritic plaques in AD cortex. Control tissue illustrates the distribution of SCAI-positive cell bodies and processes in the absence of AD pathology. Numbered regions indicate areas selected for higher-magnification imaging shown in Figure 7F–G; for example, sample AD2 region 4 for CSNK1A1 (Figure 7G), and sample AD2 region 6 for SCAI (Figure 7F).*

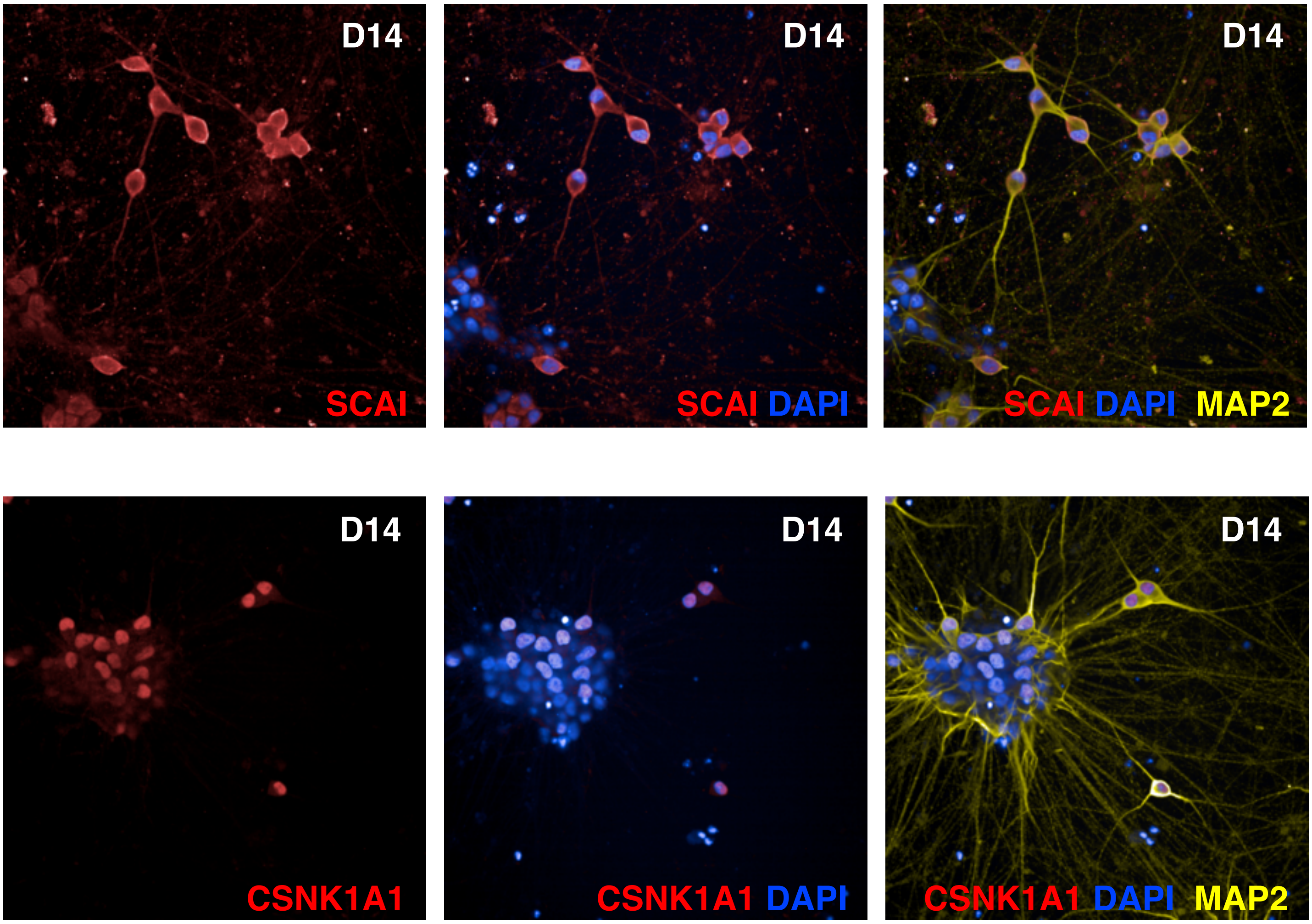


**Supplementary Figure S7. Subcellular localisation of SCAI and CSNK1A1 in cortical i3Neurons.** *Immunocytochemistry images of day-14 (D14) cortical i³Neurons stained for SCAI or CSNK1A1, together with MAP2 and DAPI. SCAI exhibits prominent staining surrounding MAP2-positive neuronal cell bodies, with weaker cytoplasmic and nuclear signal, whereas CSNK1A1 exhibits predominantly nuclear signal. Images acquired at 40× magnification.*

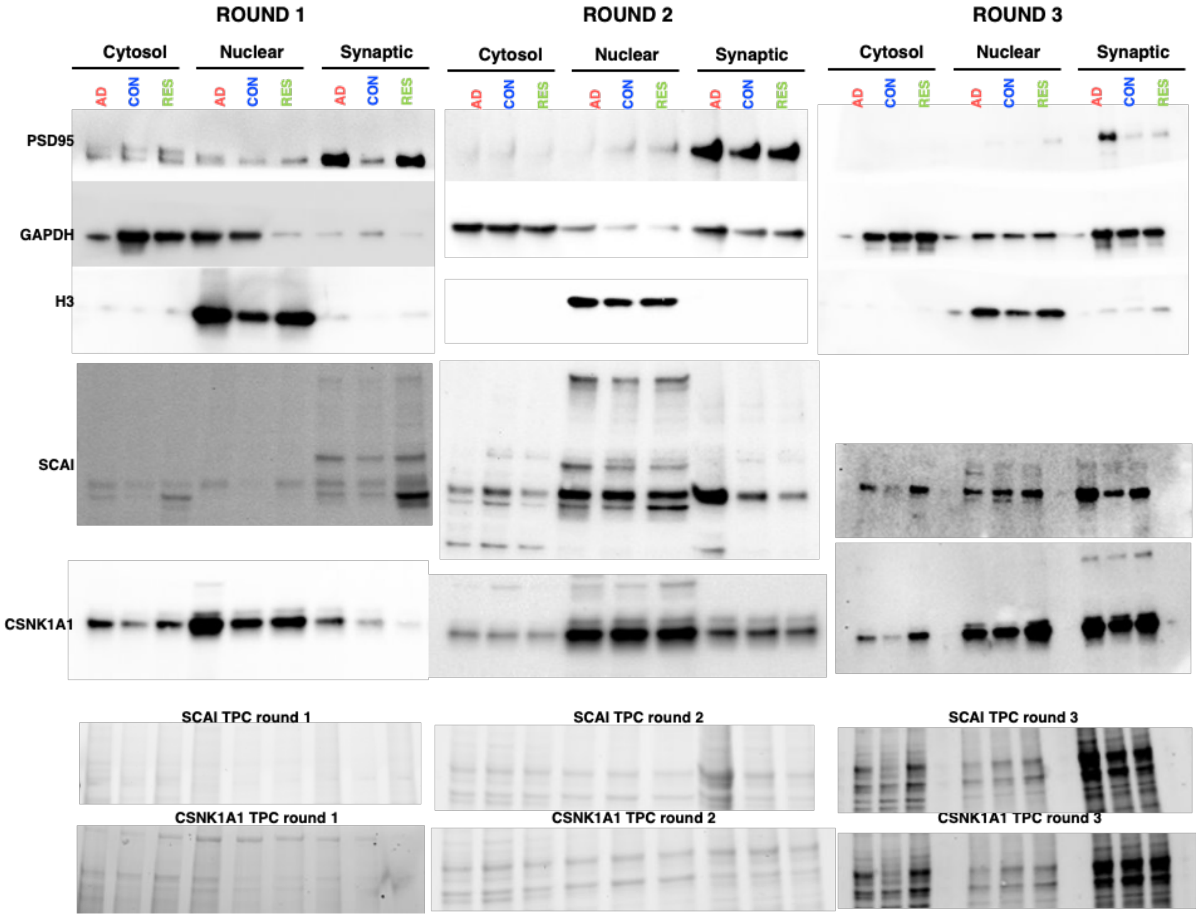


**Supplementary Figure S8. Orthogonal validation of SCAI and CSNK1A1 subcellular localisation in human dlPFC.** *Western blot analysis of cytosolic, nuclear and synaptic-enriched fractions from CON, RES and AD-DEM dlPFC (3 rounds corresponds to n = 3 subjects per condition). SCAI and CSNK1A1 localisation is shown alongside GAPDH, H3 and PSD95 as markers of cytosolic, nuclear and synaptic enrichment, respectively. Total protein content (TPC) is shown as a loading reference for SCAI and CSNK1A1.*

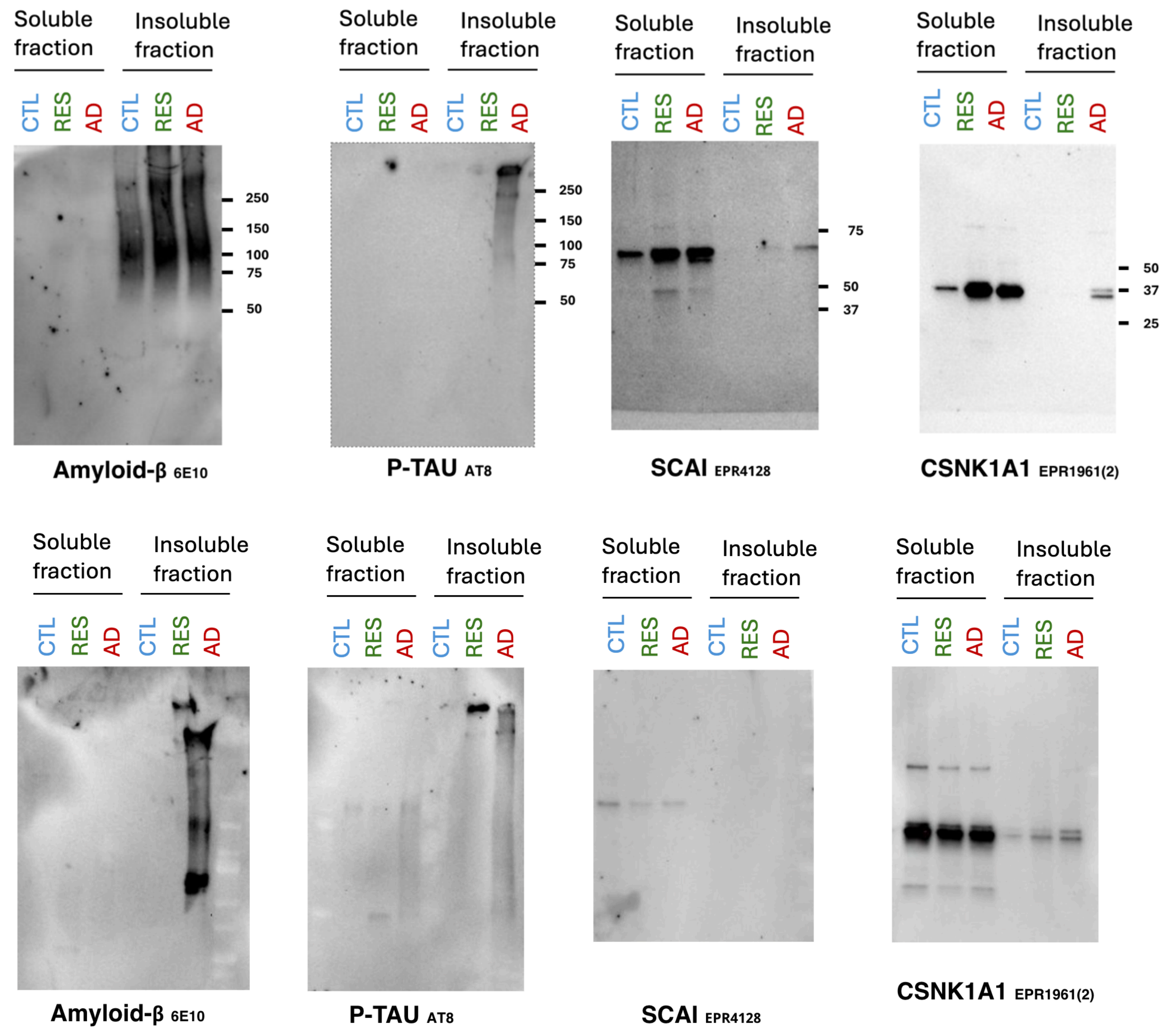


**Supplementary Figure S9. Detection of SCAI and CSNK1A1 in sarkosyl-soluble and sarkosyl-insoluble human dlPFC fractions.** *Western blot analysis of CON, RES and AD-DEM samples following sequential sarkosyl extraction (2 rounds corresponds to n = 2 subjects per condition). SCAI and CSNK1A1 are shown alongside phosphorylated tau (p-tau; AT8) and Aβ (6E10). The upper panel corresponds to the samples presented in Figure 7E; the lower panel shows the independent replicate set.*

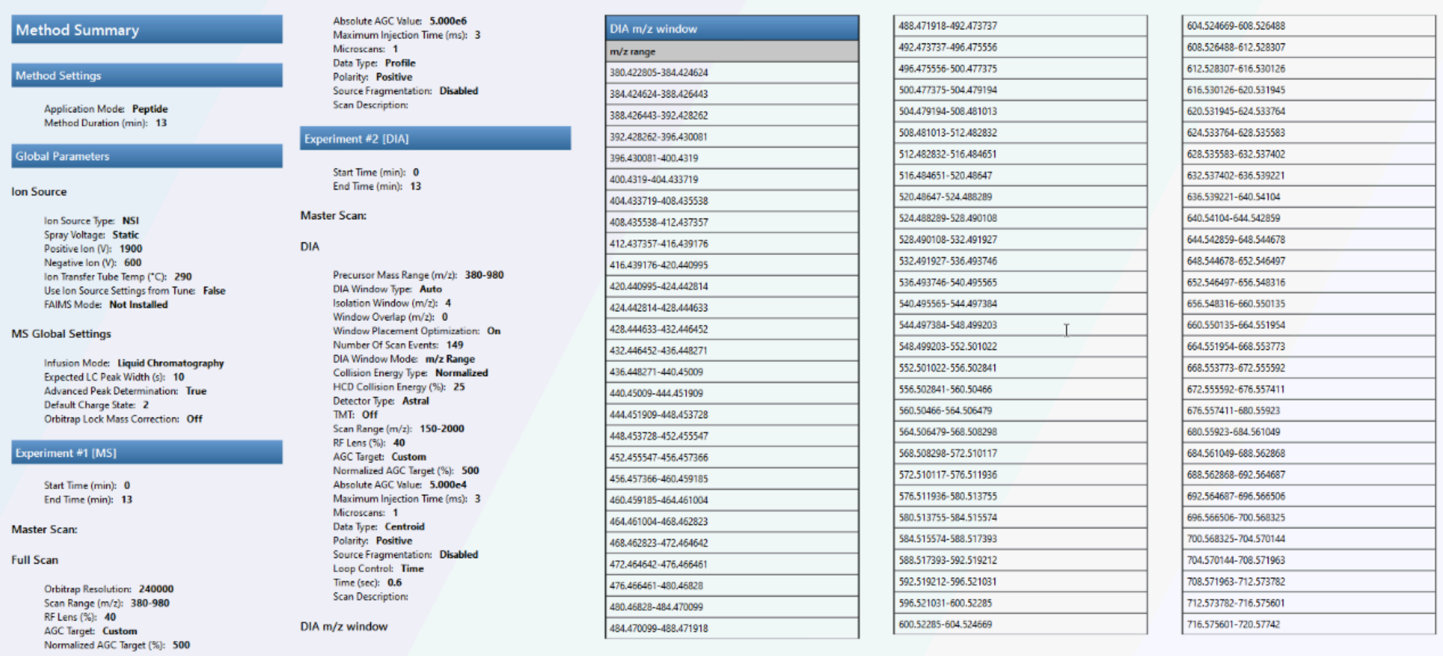

#### Method Summary

##### Method Settings

Application Mode: **Peptide**
Method Duration (min): **13**

##### Global Parameters

**Ion Source**

Ion Source Type: **NSI**
Spray Voltage: **Static**
Positive Ion (V): **1900**
Negative Ion (V): **600**
Ion Transfer Tube Temp (°C): **290**
Use Ion Source Settings from Tune: **False**
FAIMS Mode: **Not Installed**

**MS Global Settings**

Infusion Mode: **Liquid Chromatography**
Expected LC Peak Width (s): **10**
Advanced Peak Determination: **True**
Default Charge State: **2**
Orbitrap Lock Mass Correction: **Off**

##### Experiment #1 [MS]

Start Time (min): **0**
End Time (min): **13**

**Master Scan:**

**Full Scan**

Orbitrap Resolution: **240000**
Scan Range (m/z): **380-980**
RF Lens (%): **40**
AGC Target: **Custom**
Normalized AGC Target (%): **500**
Absolute AGC Value: **5.000e6**
Maximum Injection Time (ms): **3**
Microscans: **1**
Data Type: **Profile**
Polarity: **Positive**
Source Fragmentation: **Disabled**
Scan Description:

##### Experiment #2 [DIA]

Start Time (min): **0**
End Time (min): **13**

**Master Scan:**

**DIA**

Precursor Mass Range (m/z): **380-980**
DIA Window Type: **Auto**
Isolation Window (m/z): **4**
Window Overlap (m/z): **0**
Window Placement Optimization: **On**
Number Of Scan Events: **149**
DIA Window Mode: **m/z Range**
Collision Energy Type: **Normalized**
HCD Collision Energy (%): **25**
Detector Type: **Astral**
TMT: **Off**
Scan Range (m/z): **150-2000**
RF Lens (%): **40**
AGC Target: **Custom**
Normalized AGC Target (%): **500**
Absolute AGC Value: **5.000e4**
Maximum Injection Time (ms): **3**
Microscans: **1**
Data Type: **Centroid**
Polarity: **Positive**
Source Fragmentation: **Disabled**
Loop Control: **Time**
Time (sec): **0.6**
Scan Description:

DIA m/z window

| DIA m/z window |
|---|
| **m/z range** |
| 380.422805-384.424624 |
| 384.424624-388.426443 |
| 388.426443-392.428262 |
| 392.428262-396.430081 |
| 396.430081-400.4319 |
| 400.4319-404.433719 |
| 404.433719-408.435538 |
| 408.435538-412.437357 |
| 412.437357-416.439176 |
| 416.439176-420.440995 |
| 420.440995-424.442814 |
| 424.442814-428.444633 |
| 428.444633-432.446452 |
| 432.446452-436.448271 |
| 436.448271-440.45009 |
| 440.45009-444.451909 |
| 444.451909-448.453728 |
| 448.453728-452.455547 |
| 452.455547-456.457366 |
| 456.457366-460.459185 |
| 460.459185-464.461004 |
| 464.461004-468.462823 |
| 468.462823-472.464642 |
| 472.464642-476.466461 |
| 476.466461-480.46828 |
| 480.46828-484.470099 |
| 484.470099-488.471918 |

| |
|---|
| 488.471918-492.473737 |
| 492.473737-496.475556 |
| 496.475556-500.477375 |
| 500.477375-504.479194 |
| 504.479194-508.481013 |
| 508.481013-512.482832 |
| 512.482832-516.484651 |
| 516.484651-520.48647 |
| 520.48647-524.488289 |
| 524.488289-528.490108 |
| 528.490108-532.491927 |
| 532.491927-536.493746 |
| 536.493746-540.495565 |
| 540.495565-544.497384 |
| 544.497384-548.499203 |
| 548.499203-552.501022 |
| 552.501022-556.502841 |
| 556.502841-560.50466 |
| 560.50466-564.506479 |
| 564.506479-568.508298 |
| 568.508298-572.510117 |
| 572.510117-576.511936 |
| 576.511936-580.513755 |
| 580.513755-584.515574 |
| 584.515574-588.517393 |
| 588.517393-592.519212 |
| 592.519212-596.521031 |
| 596.521031-600.52285 |
| 600.52285-604.524669 |

| |
|---|
| 604.524669-608.526488 |
| 608.526488-612.528307 |
| 612.528307-616.530126 |
| 616.530126-620.531945 |
| 620.531945-624.533764 |
| 624.533764-628.535583 |
| 628.535583-632.537402 |
| 632.537402-636.539221 |
| 636.539221-640.54104 |
| 640.54104-644.542859 |
| 644.542859-648.544678 |
| 648.544678-652.546497 |
| 652.546497-656.548316 |
| 656.548316-660.550135 |
| 660.550135-664.551954 |
| 664.551954-668.553773 |
| 668.553773-672.555592 |
| 672.555592-676.557411 |
| 676.557411-680.55923 |
| 680.55923-684.561049 |
| 684.561049-688.562868 |
| 688.562868-692.564687 |
| 692.564687-696.566506 |
| 696.566506-700.568325 |
| 700.568325-704.570144 |
| 704.570144-708.571963 |
| 708.571963-712.573782 |
| 712.573782-716.575601 |
| 716.575601-720.57742 |

**Supplementary figure S10.** *Full DIA LC-MS$^2$ instrument settings.*

#### Supplementary Data:

A. Disease-group (CON/RES/AD-DEM) interaction-model results for imputed and unimputed analyses, with GO:CC enrichment of the 217 robust candidates.
B. Global-pathology interaction-model results for imputed and unimputed analyses.
C. Resilience interaction-model results for imputed and unimputed analyses.
D. Markers, TAGM-MAP predictions, OOS predictions, ranked enrichment, and JS-divergence of protein OOS predictions.
